## Supplementary information for "Coupling cellular drug-target engagement to downstream pharmacology with CeTEAM"

### Supplementary Discussion

*PARP1 L713F-GFP accumulation following proteasome and/or autophagosome inhibition*

Notably, PARP1 L713F-GFP appeared to demonstrate negligible accumulation following proteasome inhibition and detection by western blotting (**Supplementary Fig. 1g**) or live cell fluorescent microscopy (**Supplementary Fig. 1h**). Further, autophagy inhibition by bafilomycin A1 (specifically, inhibition of autophagosome-lysosome fusion) or their combination also did not affect global PARP1 L713F-GFP signal when monitored by microscopy (**Supplementary Fig. 1h**). PARP1 L713F-nLuc, however, was partially rescued by MG-132 treatment over 6 hours as seen by western blot (**Supplementary Fig. 12b**). Upon closer inspection, it became clear that proteasomal inhibition *via* MG-132 treatment resulted in specific accumulation of L713F-GFP in the nucleoli (**Supplementary Fig. 1h, and Supplementary Fig. 10c**). It has long been known that significant fractions of PARP1 reside in the nucleoli[1], but it has only recently emerged that, in addition to acting as a reservoir for the DDR, PARP1 is a key regulator of nucleolar biology[2]. Thus, this selective accumulation of L713F-GFP may relate to PARP1-associated functions in the nucleolus and will require further investigation.

*Congruence of PARP1 L713F with PARP1 biology*

PARP1 is equally amenable to GFP fusions on the N- or C-terminus without compromising functionality[3], so differential tagging of WT vs L713F PARP1 should not affect interpretation of DDR responsiveness between the two variants. Indeed, both variants were recruited to laser microirradiation-induced DNA damage sites, although L713F-GFP had slightly attenuated kinetics, as previously reported (**Supplementary Fig. 8d and e**)[4]. FRAP analyses also suggested that L713F-GFP transient nuclear mobility was slightly worse (**Supplementary Fig. 8f-j**). PARP1 WT (heterogeneous population) and L713F (clonal population) were then profiled longer-term for recruitment to damage sites and proximal markers of PARP1 activity with PARPi pre-treatment for one or 24 hours (**Supplementary Fig. 9a**, **b**, **and h**). While the mean fluorescence intensity of WT PARP1 was unaffected by PARPi alone, it temporally increased following DNA damage and then regressed to baseline (**Supplementary Fig. 9c**). However, one-hour pre-treatment with olaparib (5 μM) or talazoparib (0.5 μM) appeared to sustain higher GFP signals 60 minutes post-radiation (**Supplementary Fig. 9c and f**)[5]. PARP1 L713F mimicked WT GFP intensity in the absence of PARPi but was notably increased following one hour of PARPi treatment (**Supplementary Fig. 9b and c**). Nuclear PAR and γH2A.X trends following DNA damage were similar between PARP1 WT and L713F-expressing cells, *i.e.* PARPi suppressed PAR formation but did not affect γH2A.X (**Supplementary Fig. 9d and e**). Notably, however, basal PAR and γH2A.X levels were significantly elevated in L713F-GFP-expressing cells (**Supplementary Fig. 9b, d, and e**). PARP1 L713F-GFP remained high after 24 hours of PARPi, but both PAR and γH2A.X levels had risen even prior to DNA damage induction, which was only reciprocated in talazoparib-treated WT PARP1 cells (**Supplementary Fig. 9h-k**). This indicated that PARP trapping in the unperturbed, cycling cells had become predominant. Collectively, these data suggested that PARP1 L713F-GFP behaves similarly to PARP1 WT following PARPi treatment in the absence or presence of DNA damage.

Reintroduction of PARP1 L713F-GFP into PARP1 KO HeLa cells was previously reported to be toxic *via* NAD+ exhaustion due to continual, DNA-independent PAR formation[4]. For this reason, we utilized a doxycycline-inducible promoter in initial experiments to minimize any potential toxicity. When L713F-GFP was introduced into U-2 OS or HCT116 cells, which contain both endogenous alleles of WT PARP1, we were able to confirm elevated basal PAR and γH2A.X in comparison to WT PARP1 (**Supplementary Fig. 9d and e**) but saw no overt signs of toxicity or stress over several days/weeks of culture (**Supplementary Fig. 9g**). Nonetheless, the elevated PAR and γH2A.X levels were suppressed by PARPi treatment, as before[4] (**Supplementary Fig. 9b, d, e**). This discrepancy may reflect differences in host cell background, where the HeLa PARP1 KO cells may have physiologically adapted to PARP1 inactivation and are unable to cope with the acute, constitutive activity of PARP1 L713F. A related possibility concerns the differences in heterologous expression systems. Specifically, transient transfections can result in uneven transgene expression, where very high expression of PARP1 L713F in a fraction of target cells could become problematic for viability. The use of lentiviral transduction, where we observe better uniformity and levels of transgene expression at or below endogenous protein abundances, has likely mitigated these concerns in our experimental system.

**Supplementary Table 1. Data collection and refinement statistics of the NUDT15-NSC56456 co-crystal structure.**

| | NUDT15 + NSC56456<br>PDB ID 7NR6 |
|---|---|
| **Data collection** | |
| Beamline | BESSY 14.1 |
| Wavelength (Å) | 0.9184 |
| Space group | $P\ 2_1\ 2_1\ 2_1$ |
| Cell dimensions | |
| $a$, $b$, $c$ (Å) | 46.84, 49.03, 135.38 |
| $\alpha$, $\beta$, $\gamma$ (°) | 90, 90, 90 |
| Resolution (Å) | 67.69-1.80(1.84-1.80)* |
| $R_{merge}$ | 7.9 (91.9)* |
| *CC1/2* | 0.998 (0.757)* |
| $\langle I / \sigma I \rangle$ | 12.7 (1.9)* |
| Total observations | 199,093 (12,046)* |
| Unique observations | 29,634 (1,714)* |
| Completeness (%) | 99.7 (99.7)* |
| Redundancy | 6.7 (7.0)* |
| **Refinement** | |
| $R_{work}$ / $R_{free}$ | 18.7 / 20.6 |
| No. atoms | |
| Protein | 2502 |
| Ligand/ion | 44 |
| Water | 284 |
| Average $B$-factors (Å$^2$) | |
| Protein | 28.8 |
| Ligand/ion | 27.9 |
| Water | 38.5 |
| R.m.s. deviations | |
| Bond lengths (Å) | 0.006 |
| Bond angles (°) | 0.82 |
| Ramachandran statistics | |
| Favoured (%) | 98.7 |
| Allowed (%) | 1.3 |
| Outliers (%) | 0 |

A single crystal was used for data collection.
*Values in parentheses are for highest-resolution shell

**Supplementary Table 2.** Small molecule screening details

| Category | Parameter | Description |
|---|---|---|
| Assay | Type of assay | Cell-based, dual luminescence assay |
| | Target | PARP1 L713F (human PARP1) |
| | Primary measurement | Abundance of PARP1 L713F-nLuc relative to an akaLuc normalization signal |
| | Key reagents | • HCT116 colon carcinoma cells expressing pCW57.1-PARP1 L713F-nLuc and pLenti CMV Blast-akaLuc (see Methods)<br>• akaLumine HCl (TokeOni; Sigma Aldrich)<br>• Furimazine (Promega Nano-Glo Assay Kit) |
| | Assay protocol | See Methods section |
| | Additional comments | Luminescence readout is from intact, live cells |
| Library | Library size | 1187 |
| | Library composition | MedChemExpress Epigenetics & Selleck Nordic Oncology drug-like libraries offered by the SciLifeLab Compound Center |
| | Source | The libraries were spotted from up to 10 mM DMSO solutions in Labcyte 384 LDV plates using an Echo 550 |
| | Additional comments | SciLifeLab Compound Center spotted compounds in assay plates. |
| Screen | Format | 96-well format<br>Assay plate: Greiner Cell Culture Microplate, 96 Well, PS, F-Bottom (Chimney Well), White, Cellstar® Tc, Lid With Condensation Rings, Sterile (catalog # 655083) |
| | Concentration(s) tested | Compound concentration at 10 µM, DMSO concentration at 0.1% |
| | Plate controls | Negative control: Test cells with 0.75 µg/µL doxycycline (DOX) added + 0.1% DMSO [v/v] (Column 1 on each plate)<br>Positive control: Test cells + DOX treated with 10 µM veliparib (0.1% DMSO [v/v]; Column 2 on each plate) |
| | Reagent/ compound dispensing system | Compound dispensing system: Echo 550 from Labcyte<br>Cell dispensing system: Multidrop Combi from Thermo Scientific<br>Luminescence reagent dispensing: Eppendorf dispensing multichannel pipet |
| | Detection instrument and software | CLARIOStar microplate reader and analysis software (BMG LABTECH) |
| | Assay validation/QC | • Negative control – average nLuc/akaLuc ratio: 0.014, standard deviation: 0.004; positive control – average nLuc/akaLuc: 0.077, standard deviation: 0.018; average Z' factor/plate: 0.29.<br>• To account for extreme viability and expression variabilities, compounds with akaLuc signals deviating >4 SDs from control means on each plate were excluded from analysis.<br>• QC also included monitoring of plate edge effects and hit distribution of the hits, with no additional corrections necessary. |
| | Correction factors | Exclusion of compounds resulting in akaLuc signals >4 SDs from controls per plate. |
| | Normalization | L713F-nLuc signal is normalized to the respective akaLuc signal in each well (nLuc/akaLuc ratio). For simplified data visualization, the nLuc/akaLuc ratio was set relative to the mean for the negative control (+DOX, DMSO) to generate fold-change. |
| | Additional comments | None |
| Post-HTS analysis | Hit criteria | Hit threshold: Average $log_2$(nLuc/akaLuc) ratio of test samples with akaLuc <4 SDs (-0.0148) ± 2xSD = -1.5501 (negative) or 1.4304 (positive). Hits were also defined as +3xSDs = 2.2881. |
| | Hit rate | 6.31% (53/840) |
| | Additional assay(s) | Hit confirmation was performed with the same assay conditions (10 µM final concentration) in triplicate with 1-2 sets (total of up to 6 replicates) |
| | Confirmation of hit purity and structure | Hits confirmed by LCMS |
| | Additional comments | None |

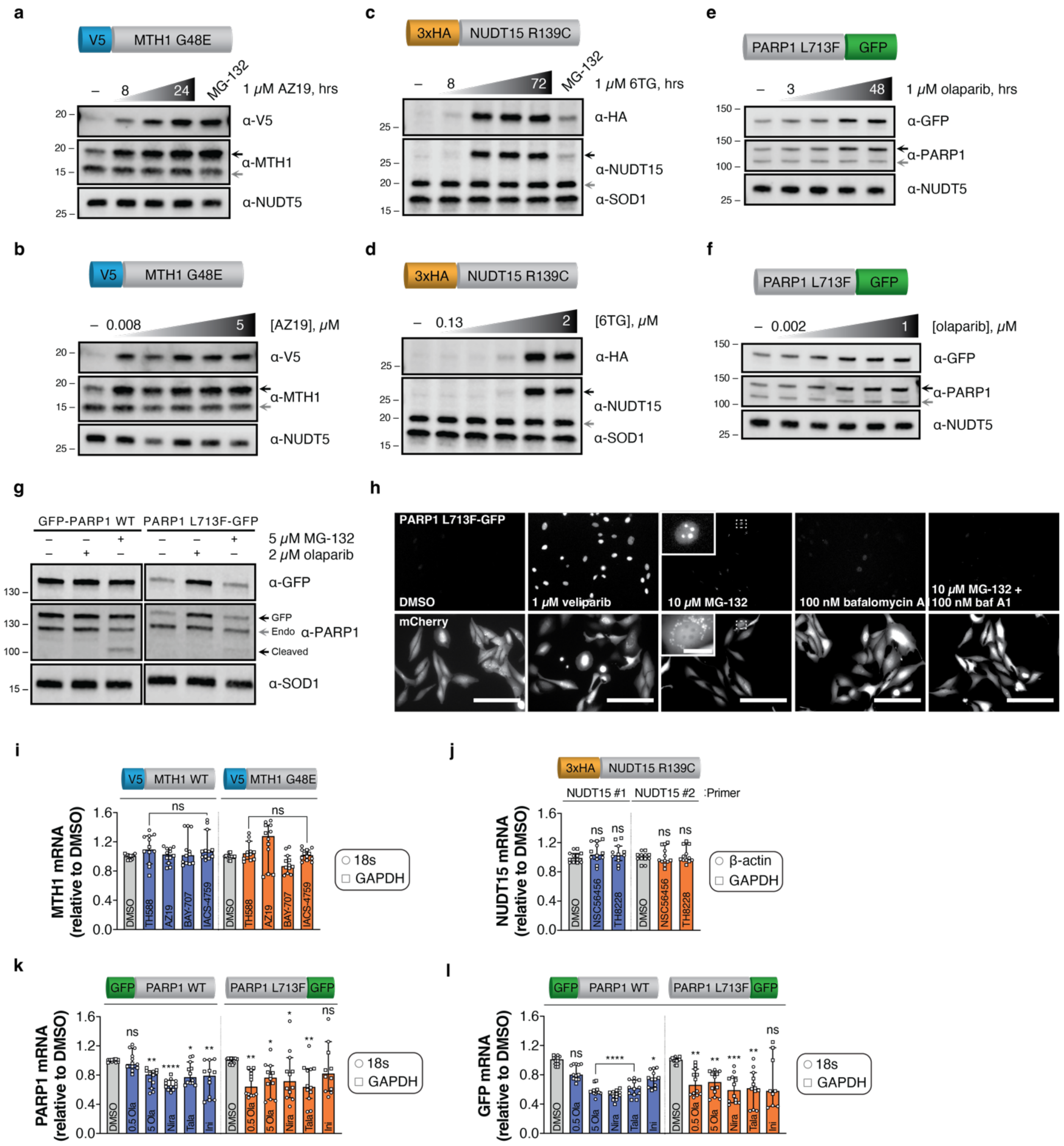
a
V5 MTH1 G48E
– 8 24 MG-132
1 μM AZ19, hrs
α-V5
α-MTH1
α-NUDT5
b
V5 MTH1 G48E
– 0.008 5
[AZ19], μM
α-V5
α-MTH1
α-NUDT5
c
3xHA NUDT15 R139C
– 8 72 MG-132
1 μM 6TG, hrs
α-HA
α-NUDT15
α-SOD1
d
3xHA NUDT15 R139C
– 0.13 2
[6TG], μM
α-HA
α-NUDT15
α-SOD1
e
PARP1 L713F GFP
– 3 48
1 μM olaparib, hrs
α-GFP
α-PARP1
α-NUDT5
f
PARP1 L713F GFP
– 0.002 1
[olaparib], μM
α-GFP
α-PARP1
α-NUDT5
g
GFP-PARP1 WT
PARP1 L713F-GFP
5 μM MG-132
2 μM olaparib
α-GFP
GFP
Endo
Cleaved
α-PARP1
α-SOD1
h
PARP1 L713F-GFP
DMSO
1 μM veliparib
10 μM MG-132
100 nM bafalomycin A
10 μM MG-132 + 100 nM baf A1
mCherry
i
V5 MTH1 WT
V5 MTH1 G48E
MTH1 mRNA (relative to DMSO)
DMSO
TH588
AZ19
BAY-707
IACS-4759
ns
18s
GAPDH
j
3xHA NUDT15 R139C
NUDT15 #1
NUDT15 #2
:Primer
NUDT15 mRNA (relative to DMSO)
DMSO
NSC56456
TH8228
ns
β-actin
GAPDH
k
GFP PARP1 WT
PARP1 L713F GFP
PARP1 mRNA (relative to DMSO)
DMSO
0.5 Ola
5 Ola
Nira
Tala
Ini
18s
GAPDH
l
GFP PARP1 WT
PARP1 L713F GFP
GFP mRNA (relative to DMSO)
DMSO
0.5 Ola
5 Ola
Nira
Tala
Ini
18s
GAPDH

**Supplementary Figure 1. Initial time and dose characterization of MTH1, NUDT15, and PARP1 drug biosensors (related to Figure 1). a**, Representative blot (n=2) of U-2 OS V5-MTH1 G48E cells incubated with DMSO (24 hours), 5 μM MG-132 (6 hours), or 1 μM AZ19 (8-24 hours). **b**, Representative blot (n=2) of U-2 OS V5-MTH1 G48E cells incubated with DMSO or AZ19 (0.008-5 μM) for 24 hours. **c**, Representative blot (n=2) of HCT116 3-6 3xHA-NUDT15 R139C cells incubated with DMSO (72 hours), 5 μM MG-132 (6 hours), or 1 μM 6TG (8-72 hours). **d**, Representative blot (n=2) of HCT116 3-6 3xHA-NUDT15 R139C cells incubated with DMSO or 6TG (0.13-2 μM for 72 hours. **e**, Representative blot (n=2) of U-2 OS PARP1 L713F-GFP cells incubated with DMSO (48 hours) or 1 μM olaparib (3-48 hours). **f**, Representative blot (n=2) of U-2 OS PARP1 L713F-GFP cells incubated with DMSO or olaparib (0.0016-1 μM) for 48 hours. **g**, Representative blot (n=2) of U-2 OS GFP-PARP1 WT or L713F-GFP cells incubated with DMSO, 2 μM olaparib (24 hours), or 5 μM MG-132 (6 hours) and prepared for western blot with the indicated antibodies. Black arrow – mutant fusion protein; grey arrow – endogenous protein. **h**, U-2 OS PARP1 L713F-GFP/mCherry cells were incubated with DMSO, 1 μM veliparib (24 hours), 10 μM MG-132, 100 nM bafilomycin A1, or 10 μM MG-132 and 100 nM bafilomycin A1 (each for 6 hours) before live cell fluorescence microscopy. Scale bar is 200 μm and 20 μm for inset. **i**, U-2 OS V5-MTH1 WT or G48E cells were treated with DMSO, 2.5 μM TH588, 1 μM AZ19, 1 μM BAY-707, or 1 μM IACS-4759 for 24 hours before RT-qPCR analysis. **j**, HCT116 3-6 3xHA-NUDT15 R139C cells were treated with DMSO, 20 μM NSC56456, or 20 μM TH8228 for 24 hours before RT-qPCR analysis. **k**, U-2 OS GFP-PARP1 WT or PARP1 L713F-GFP cells were treated with DMSO, 0.5 or 5 μM olaparib, 5 μM niraparib, 0.5 μM talazoparib, or 10 μM iniparib for 24 hours before RT-qPCR analysis. Total PARP1 transcripts or GFP transcripts (**l**) were quantified. Median values from two independent experiments run in triplicate and normalized to each housekeeping gene with 95% confidence intervals plotted. In all cases, ns – not significant; * – $p < 0.05$; ** – $p < 0.01$; *** – $p < 0.001$; **** – $p < 0.0001$ by Kruskal-Wallis test with multiple comparisons to DMSO controls for each primer or cell set (Dunn's test; Kruskal-Wallis statistic = $4.457_{\textbf{MTH1 WT}}$, $13.23_{\textbf{G48E}}$, $1.665_{\textbf{NUDT15}}$, $0.7138_{\textbf{R139C}}$, $35.00_{\textbf{PARP1 WT-PARP1}}$, $18.07_{\textbf{L713F-PARP1}}$, $55.01_{\textbf{PARP1 WT-GFP}}$, $24.03_{\textbf{L713F-GFP}}$). In all cases, cells were induced with doxycycline before drug addition.

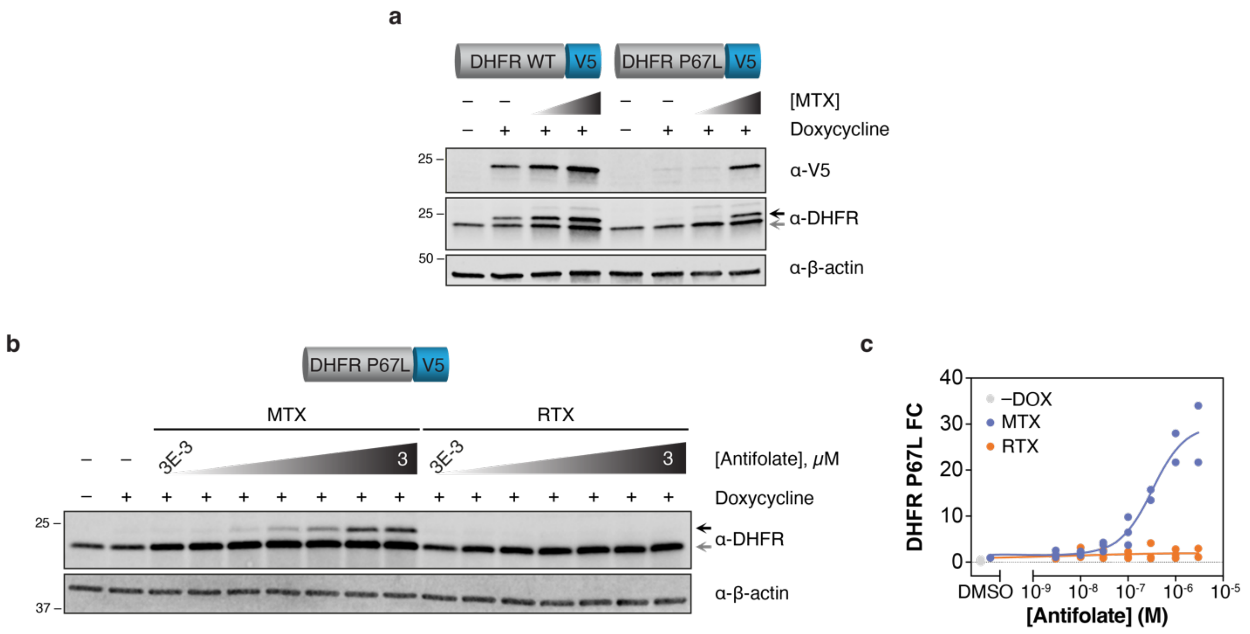

**Supplementary Figure 2. DHFR P67L detects intracellular DHFR binders. a**, A representative western blot (n=2) comparing WT and P67L DHFR in response to methotrexate. DHFR WT-V5 and DHFR P67L-V5 expression was induced in U-2 OS cells for 24 hours with doxycycline before addition of DMSO, 10 nM, or 1 μM methotrexate (MTX) for 24 hours. Black arrow – exogenous fusion protein; grey arrow – endogenous protein. **b**, A representative western blot (n=3) of DHFR P67L-V5 stabilization by methotrexate (MTX) or raltitrexed (RTX). DHFR P67L-V5 was induced with doxycycline for 24 hours, then incubated with the indicated concentration of anti-folate. Black arrow – exogenous fusion protein; grey arrow – endogenous protein. **c**, Summary quantification of DHFR P67L-V5 abundance following MTX (blue) or RTX (orange) concentration gradient. A no (–) DOX DMSO control (grey) is included for reference. Individual data points for three independent experiments and respective curve fitting are shown.

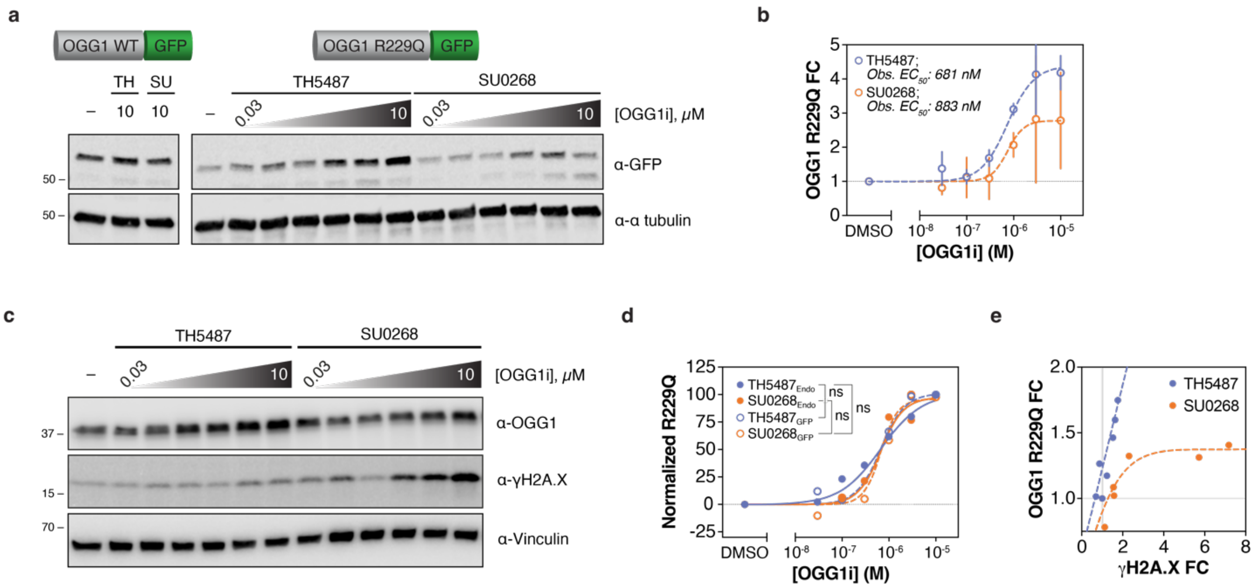

**Supplementary Figure 3. Endogenous and exogenous OGG1 R229Q as a biosensor of OGG1 inhibitor binding. a**, A representative western blot (n=3) of OGG1 WT-GFP or OGG1 R229Q-GFP responses to TH5487 or SU0268. WT or R229Q-GFP-expressing U-2 OS cells were incubated with DMSO or OGG1i for 24 hours. **b**, Summary quantification of OGG1 R229Q-GFP abundance following TH5487 (blue) or SU0268 (orange) concentration gradient. Means from three independent experiments ± SD and respective curve fittings are shown. **c**, A representative western blot (n=3) of endogenous OGG1 R229Q stabilization by OGG1i. KG-1 cells were incubated with DMSO or OGG1i for 24 hours. **d**, Overlay of exogenous (open circles) and endogenous (closed circles) OGG1 R229Q following a TH5487 (blue) or SU0268 (orange) concentration gradient normalized to the highest dataset value. Means from three independent experiments are shown with respective curve fittings. ns – not significant by extra sum-of-squares F Test (*F*: 0.4290, DFn: 3, DFd: 73)$_{\text{all}}$, (*F*: 0.5406, DFn: 1, DFd: 33)$_{\text{SU0268}}$, (*F*: 0.1861, DFn: 1, DFd: 32)$_{\text{TH5487}}$. **e**, Two-dimensional summary of R229Q stabilization and γH2A.X induction following TH5487 (blue) or SU0268 (orange) concentration gradients. Means of three independent experiments are shown along with linear (TH5487, Y = 0.5863*X + 0.5846) or non-linear (logistic growth [$r^2$:0.4464]; SU0268) lines-of-best-fit. FC – fold change.

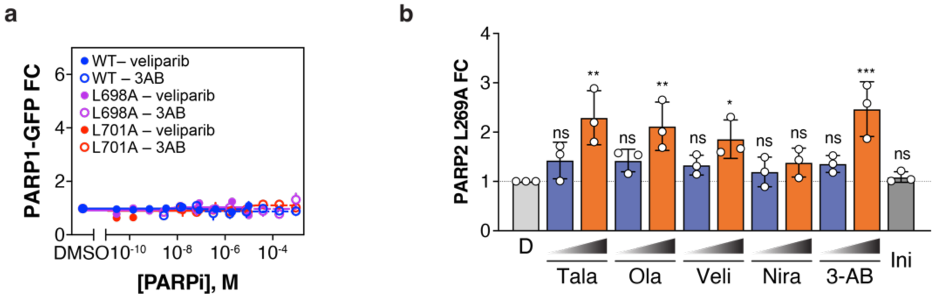

**Supplementary Figure 4. Rational expansion of CeTEAM-capable mutations from PARP1 L713F (related to Figure 2). a**, Live-cell fluorescence fold change of WT (blue), L698A (purple), or L701A (red) PARP1-GFP CeTEAM variants with veliparib (closed circles) or 3-AB (open circles) dose-response after 24 hours. Means of n=2 experiments shown ± SEM. **b**, Summary western blot quantitation data of PARP2 L269A-GFP fold change following low (blue) or high (orange) concentration incubation of talazoparib, olaparib, veliparib, niraparib (10 nM/1 µM), or 3-aminobenzamide (3-AB, 10 µM/1 mM) incubation for 24 hours and relative to DMSO control (light grey). 20 µM iniparib is included as a negative control (dark grey), and the mean of three independent experiments is shown with SD denoted. ns – not significant, * – $p < 0.05$, ** – $p < 0.01$, *** – $p < 0.001$ by ordinary one-way ANOVA with multiple comparisons to the DMSO control (Dunnett's test; $F_{Treatment}$ [DFn, DFd] = 5.842 [11, 24]). RFU – relative fluorescence units, FC – fold change.

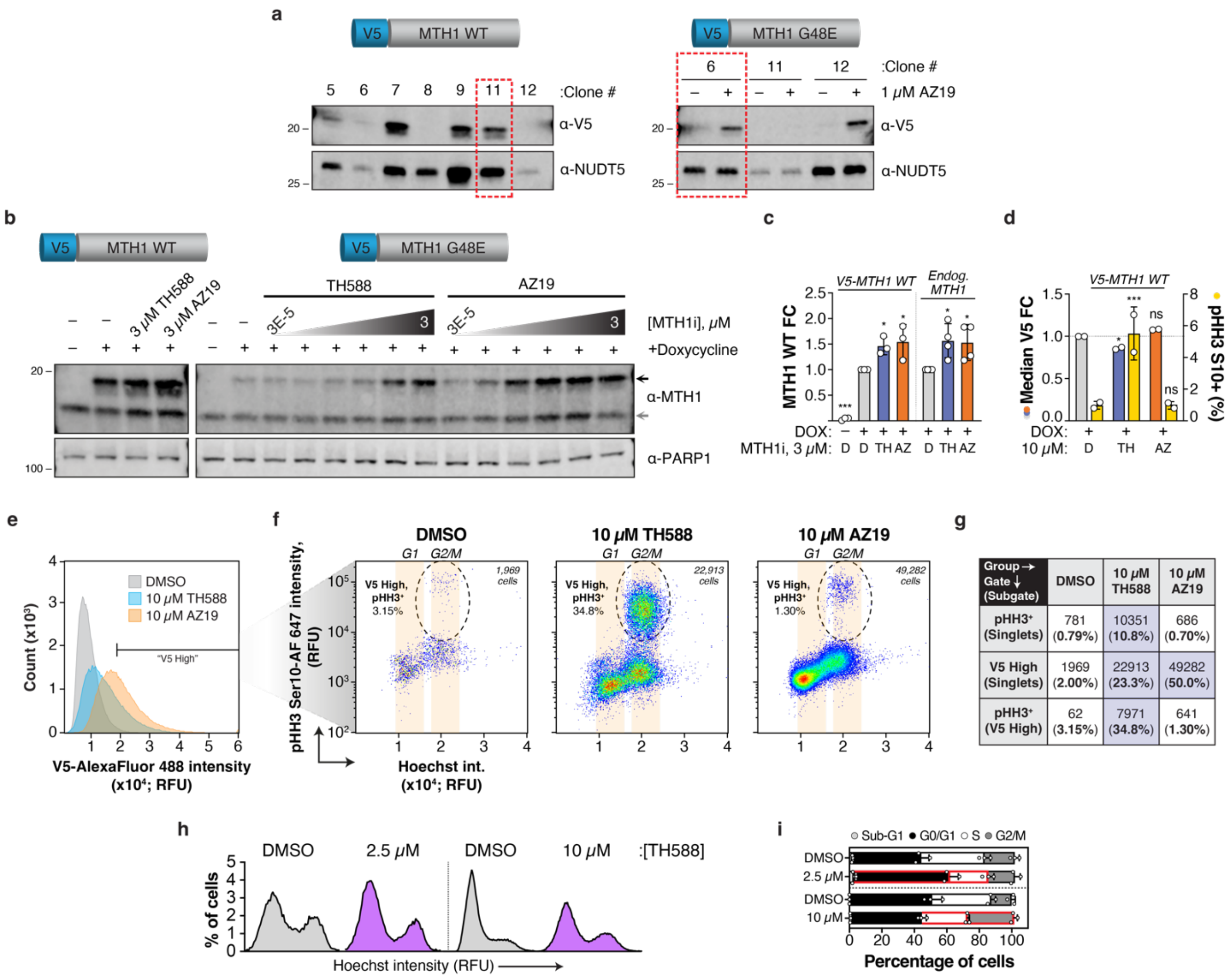

**Supplementary Figure 5. Optimization and characterization of the MTH1 G48E drug biosensor (related to Figure 3).** **a**, Clonal selection of WT (clone 11) and G48E (clone 6) MTH1 cells. Single cell clones were obtained by serial dilution into 96-well plates and testing of transgene expression (WT) or expression and accumulation dynamic range following MTH1i (G48E). **b**, A representative blot (n=3) of MTH1 G48E saturation curves generated with either TH588 or AZ19 after 24-hour incubation. MTH1 WT cells were incubated with 3 µM of either drug for comparison. Transgene expression was induced by 24-hour pre-incubation with doxycycline. Black arrow – exogenous fusion protein; grey arrow – endogenous protein. **c**, Summary western blot quantitation of exogenous (left) or endogenous (right) MTH1 WT following exposure to DMSO (light grey), 3 µM TH588 (blue), or 3 µM AZ19 (orange) for 24 hours. No (–) DOX control is included for reference. Means of three (exogenous) or four (endogenous) independent experiments ± SD are shown. **d**, Summary quantitation of V5-MTH1 WT (fold change, left axis) and pHH3 Ser10 (%, right axis, yellow) following 24-hour MTH1i treatment at 10 µM by flow cytometry. Means of two independent experiments ± SD are shown. For **c** and **d**, ns – not significant, * – $p < 0.05$, *** – $p < 0.001$ by ordinary one-way ANOVA with multiple comparisons to the DMSO control (Dunnett's test; $F_{Treatment\ (\mathbf{c\ left})}$ [DFn, DFd] = 39.86 [3, 8], $F_{Treatment\ (\mathbf{c\ right})}$ [DFn, DFd] = 5.185 [2, 9], $F_{Treatment\ (\mathbf{d\ V5})}$ [DFn, DFd] = 25.58 [10, 11], $F_{Treatment\ (\mathbf{d\ HH3})}$ [DFn, DFd] = 49.67 [10, 21]). **e**,

Representative histograms of V5-G48E intensity and "V5 high" events – arbitrarily classified as the top 2% of DMSO control signal intensity – following 24-hour TH588 (blue) or AZ19 (orange) treatment at 10 μM. **f**, "V5 high" cells were plotted by pHH3 Ser10 (y-axis) and Hoechst intensity (x-axis). "V5 high" and pHH3+ cells are identified by dashed circles and the percentages of pHH3+ cells are shown. **g**, Number of positive events and proportion of parental population (in parentheses, %) in specific sub-populations. Statistics of note are highlighted in blue. **e**/**f**/**g** show a representative experiment from n=3. **h**, Representative (from n=3) cell cycle histograms (Hoechst intensity) comparing 2.5 and 10 μM TH588 (purple) to respective DMSO (grey) controls after 24 hours. DMSO (2.5 μM)=19706 cells, 2.5 μM TH588=19429, DMSO (10 μM)=99817 cells, and 10 μM TH588=99623 cells. **i**, Cell cycle proportions by Hoechst intensity, related to **h**. Mean of three independent experiments ± SD. The 10 μM dataset is also presented in **Figure 3f**. Red highlight – adjusted $p < 0.05$ by multiple unpaired t test with Welch's correction (**2.5 μM**; t ratio/df = $1.193/2.215_{\text{Sub-G1}}$, $4.257/3.980_{\text{G1/G0}}$, $5.466/4.000_{\text{S}}$, $0.9586/3.914_{\text{G2/M}}$; **10 μM**; t ratio/df = $7.069/2.003_{\text{Sub-G1}}$, $2.119/2.961_{\text{G1/G0}}$, $5.324/2.823_{\text{S}}$, $9.956/3.994_{\text{G2/M}}$)

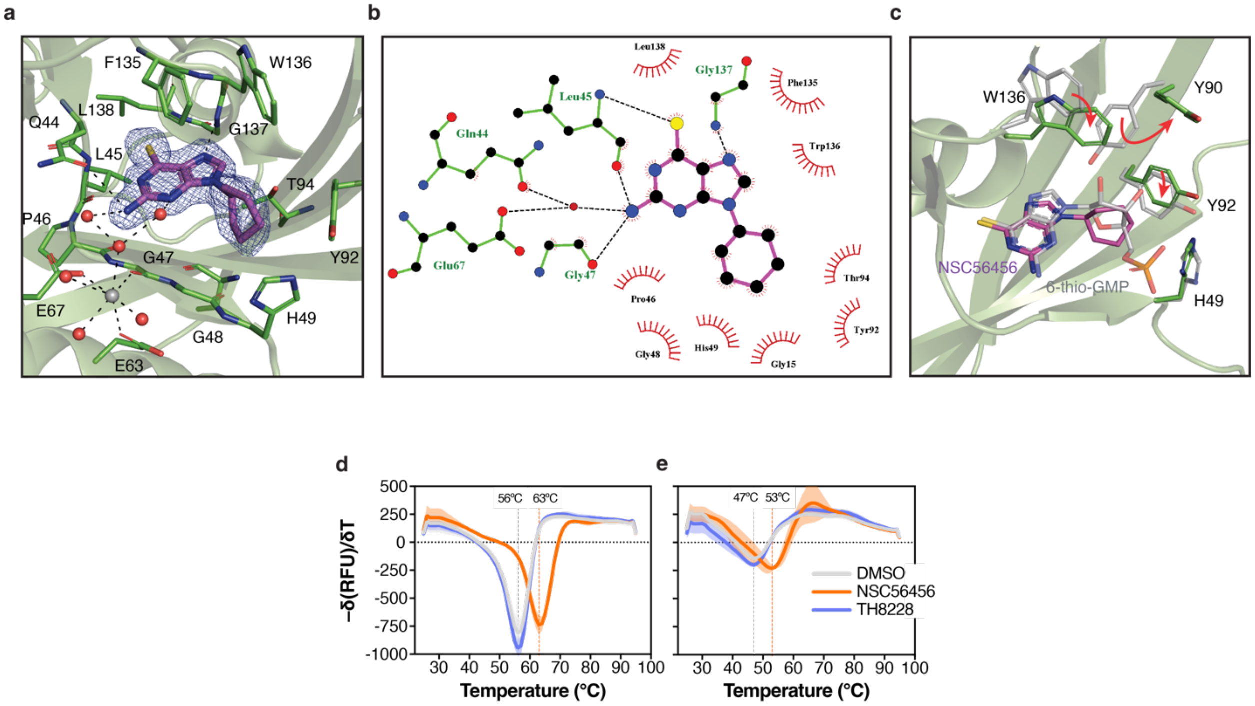

**Supplementary Figure 6. NSC56456, but not the inactive analog, TH8228, binds to NUDT15 WT and the R139C missense variant (related to Figure 4). a**, Co-crystal structure of NSC56456 bound to the catalytic site of WT NUDT15 with binding interactions. NSC56456 is depicted in magenta (with $2F_o$-$F_c$ electron density map in blue), interacting NUDT15 residues are in bright green, magnesium ions in grey, and water molecules in red. Hydrogen bonding interactions are depicted by dashed, black lines. **b**, Ligplot+ representation of NSC56456 binding interactions in the NUDT15 active site. Residues interacting with NSC56456 (magenta) are shown in green. Hydrophobic interactions are shown as an arc with spokes and hydrogen bonds as dashed lines. **c**, Binding modality comparison of NSC56456 and 6-thio-GMP within NUDT15. The aligned 6-thio-GMP-bound structure of NUDT15 (PDB ID: 5LPG) is shown in grey, while the NUDT15-NSC56456 structure is green. Residues that undergo a significant conformational change between the two structures are highlighted. The 6-thioguanine group is positioned in nearly the exact same position in both structures, aside from a slight tilt. This tilt strengthens several hydrogen bonds by bringing the bonding partners into closer proximity (bonding to Leu45 is decreased by 0.1 Å and Gly137 by 0.4 Å, respectively, **b**). The cyclohexane ring is situated similarly to ribose in the 6-thio-GMP structure, but since it lacks the protruding hydroxyl groups, this leaves empty space in the binding pocket. Trp136 has moved into the active site by 2.9 Å to fill this empty space, which causes Tyr90 to be flipped outwards. **d**, DSF of NUDT15 WT with 50 µM NSC56456 (orange) or TH8228 (blue) compared to DMSO (grey) and displayed as the negative first derivative. Means of n=2 independent experiments ± SD (shading) shown. **e**, DSF of NUDT15 R139C with 50 µM NSC56456 (orange) or TH8228 (blue) compared to DMSO (grey) and displayed as the negative first derivative. Means of n=2 independent experiments ± SD (shading) shown. **d** and **e** are related to **Figure 4e**.

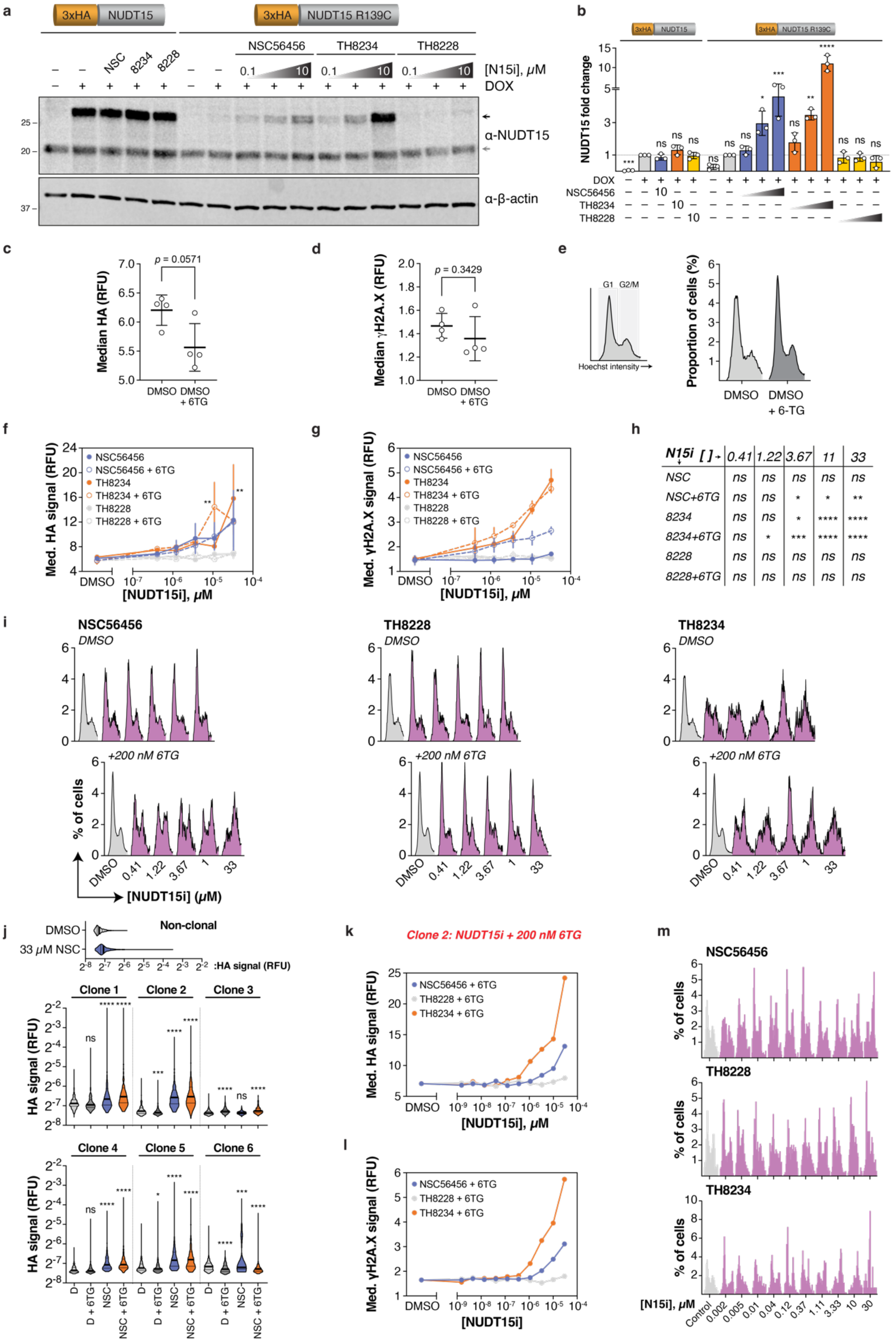
a
3xHA NUDT15
3xHA NUDT15 R139C
NSC56456
TH8234
TH8228
NSC
8234
8228
0.1 10
[N15i], μM
DOX
α-NUDT15
α-β-actin
b
NUDT15 fold change
DOX
NSC56456
TH8234
TH8228
c
Median HA (RFU)
p = 0.0571
DMSO
DMSO + 6TG
d
Median γH2A.X (RFU)
p = 0.3429
e
G1 G2/M
Hoechst intensity
Proportion of cells (%)
DMSO
DMSO + 6-TG
f
Med. HA signal (RFU)
NSC56456
NSC56456 + 6TG
TH8234
TH8234 + 6TG
TH8228
TH8228 + 6TG
[NUDT15i], μM
g
Med. γH2A.X signal (RFU)
h
N15i [ ] 0.41 1.22 3.67 11 33
NSC ns ns ns ns ns
NSC+6TG ns ns * * **
8234 ns ns * **** ****
8234+6TG ns * *** **** ****
8228 ns ns ns ns ns
8228+6TG ns ns ns ns ns
i
NSC56456
TH8228
TH8234
DMSO
+200 nM 6TG
% of cells
[NUDT15i] (μM)
j
Non-clonal
DMSO
33 μM NSC
:HA signal (RFU)
Clone 1
Clone 2
Clone 3
Clone 4
Clone 5
Clone 6
HA signal (RFU)
D
D + 6TG
NSC
NSC + 6TG
k
Clone 2: NUDT15i + 200 nM 6TG
Med. HA signal (RFU)
NSC56456 + 6TG
TH8228 + 6TG
TH8234 + 6TG
[NUDT15i], μM
l
Med. γH2A.X signal (RFU)
[NUDT15i]
m
NSC56456
TH8228
TH8234
% of cells
[N15i], μM
Control

**Supplementary Figure 7. Establishing and optimizing NUDT15 R139C as a thiopurine biosensor (related to Figure 4).** **a**, A representative western blot of NUDT15 WT and R139C accumulation following 24-hour incubation with prospective thiopurine-mimetic NUDT15i at the indicated concentrations (0.1, 1, or 10 µM). NUDT15 WT cells were incubated with 10 µM of NUDT15i for comparison. Black arrow – exogenous fusion protein; grey arrow – endogenous protein. **b**, Summary quantification of 3xHA-NUDT15 WT or R139C abundance following NSC56456 (blue), TH8234 (orange), or TH8228 (yellow) concentration gradient and compared to DMSO (light grey). No (–) DOX control (dark grey) is included for reference. Means from three independent experiments ± SD are shown. **c**, Median HA and (**d**) γH2A.X intensities of individual HCT116 3-6 3xHA-NUDT15 R139C cells treated with DMSO or 200 nM 6TG for 72 hours. Two independent experiments performed in duplicate. **e**, Representative Hoechst histograms related to **c** and **d**. Cell cycle profiles inferred from $n_{DMSO}$: 25,371 cells or $n_{6TG}$: 47,435 cells. **f**, Summary median HA or γH2A.X (**g**) intensity of induced HA-R139C cells pre-incubated with NSC56456 (blue), TH8234 (orange), or TH8228 (grey) for 3 hours followed by DMSO (closed circles) or 200 nM 6TG (open circles) for an additional 72 hours. Means of two independent experiments and ranges are shown. Representative of 1000 cells per sample. **h**, Statistical significance overview of **g** (NUDT15i x dose). **i**, Representative cell cycle profiles (Hoechst histograms) of cell populations treated as above (n=2). See Source Data for cell numbers analyzed. **j**, HA signal intensity violin plots of induced 3xHA-NUDT15 R139C clonal cells treated with DMSO or 30 µM NSC56456 -/+ 200 nM 6TG for 72 hours. n=500 cells per sample. Non-clonal cells are included as an inset for comparison. Medians (thick lines) and quartiles (thin lines) are shown on each plot. **k**, Representative data of median HA and γH2A.X (**l**) signal from 500 representative cells following pre-incubation of induced clone 2 cells with NSC56456 (blue), TH8234 (orange), or TH8228 (grey) followed by 200 nM 6TG. **m**, Cell cycle profiles (Hoechst histograms) for clone 2 cells (related to **k** and **l**). See Source Data for cell numbers analyzed. In all cases, ns – not significant, * – $p < 0.05$, ** – $p < 0.01$, *** – $p < 0.001$, and **** – $p < 0.0001$ by ordinary one-way ANOVA with multiple comparisons to the WT DMSO control (Dunnett's test; $F_{Treatment}$ [DFn, DFd] = 48.46 [15, 32]; **b**), ordinary two-way ANOVA with multiple comparisons to the DMSO control (Dunnett's test; **f**, **g**/**h**; F [DFn, DFd]: $F_{Interaction\ \mathbf{f}}$ [25, 35] = 0.8319, $F_{Row\ Factor\ \mathbf{f}}$ [5, 35] = 6.161, $F_{Column\ Factor\ \mathbf{f}}$ [5, 35] = 3.379; F [DFn, DFd]: $F_{Interaction\ \mathbf{g/h}}$ [25, 36] = 6.195, $F_{Row\ Factor\ \mathbf{g/h}}$ [5, 36] = 26.45, $F_{Column\ Factor\ \mathbf{g/h}}$ [5, 36] = 45.94), Kruskal-Wallis test with comparison to the DMSO control for each set (Dunn's test; **j**; Kruskal-Wallis statistic = $278.7_{\mathbf{clone1}}$, $1134_{\mathbf{clone2}}$, $299.3_{\mathbf{clone3}}$, $730.4_{\mathbf{clone4}}$, $536.2_{\mathbf{clone5}}$, $168.5_{\mathbf{clone6}}$), or two-tailed Mann-Whitney U test (**c** and **d**; Mann-Whitney U = $1_{\mathbf{c}}$, $4_{\mathbf{d}}$). RFU – relative fluorescence units, purple histogram – treated with NUDT15i.

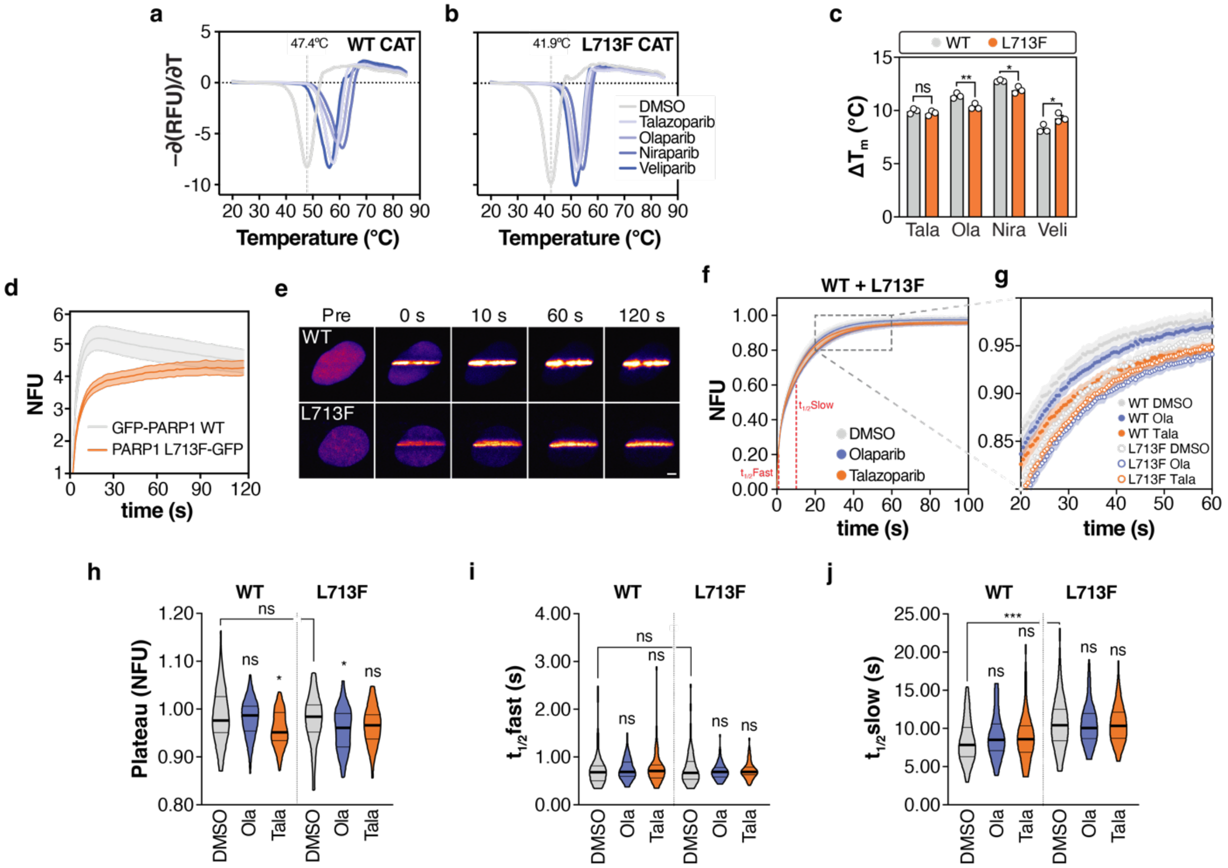

**Supplementary Figure 8. Biophysical characterization of PARPi binding to, DNA damage recruitment of, and mobility of PARP1 L713F. a**, Representative DSF with 5 µM PARP1 WT or (**b**) L713F catalytic domain with DMSO or 250 µM PARPi displayed as the negative first derivative. Melting temperatures ($T_m$) of DMSO controls are shown from duplicate readings. **c**, $T_m$ change with PARPi relative to DMSO ($\Delta T_m$). Means of three independent experiments ± SD shown. **d**, Quantification of live-cell recruitment kinetics for GFP-PARP1 WT and PARP1 L713F-GFP to laser microirradiation-induced DNA damage sites. Mean values ± SEM are shown from two independent experiments ($n_{WT}$: 20 total cells, $n_{L713F}$: 19 total cells). **e**, Representative snapshots of PARP1 saturation at microirradiation-induced DNA breaks from **d**. Scale bar = 5 µm. **f**, FRAP profiles of PARP1 WT and L713F-GFP with PARPi. GFP-PARP1 WT or L713F-GFP-expressing U-2 OS cells were pre-treated with DMSO, 5 µM olaparib, or 0.5 µM talazoparib for 24 hours prior to FRAP experiments. Approximate half-time for fast and slow components of FRAP curve are labeled. $n_{WT\text{-}DMSO} = 45$, $n_{L713F\text{-}tala} = 49$; all others n = 50 compiled from five independent experiments. **g**, Zoomed inset from **f**. **h**, Violin plots of the plateau, fast half-time ($t_{1/2}$fast; **i**), and slow half-time ($t_{1/2}$slow; **j**) from FRAP experiments of WT or L713F cells treated with DMSO (grey), olaparib (blue), or talazoparib (orange). ns – not significant; * – $p < 0.05$; ** – $p < 0.01$, *** – $p < 0.001$ by multiple unpaired t test with Welch's correction (**b;** t ratio/df = $1.017/3.991_{tala}$, $4.641/3.977_{ola}$, $4.444/2.297_{nira}$, $3.131/3.946_{veli}$), Kruskal-Wallis analyses were performed with comparisons to DMSO treatments for each group (Dunn's test; **h**, **i**, and **j**; Kruskal-Wallis statistic = $10.87_{\mathbf{h,WT}}$, $5.713_{\mathbf{h,L713F}}$, $0.8287_{\mathbf{i,WT}}$, $0.8899_{\mathbf{i,L713F}}$, $1.218_{\mathbf{j,WT}}$, $0.08776_{\mathbf{j,L713F}}$), or two-tailed Mann-Whitney U test (**h**, **i**, and **j** [comparison of WT and L713F DMSO controls]; Mann-Whitney U = $1088_{\mathbf{h}}$, $1095_{\mathbf{i}}$, $664_{\mathbf{j}}$). NFU – normalized fluorescence units, RFU – relative fluorescence units.

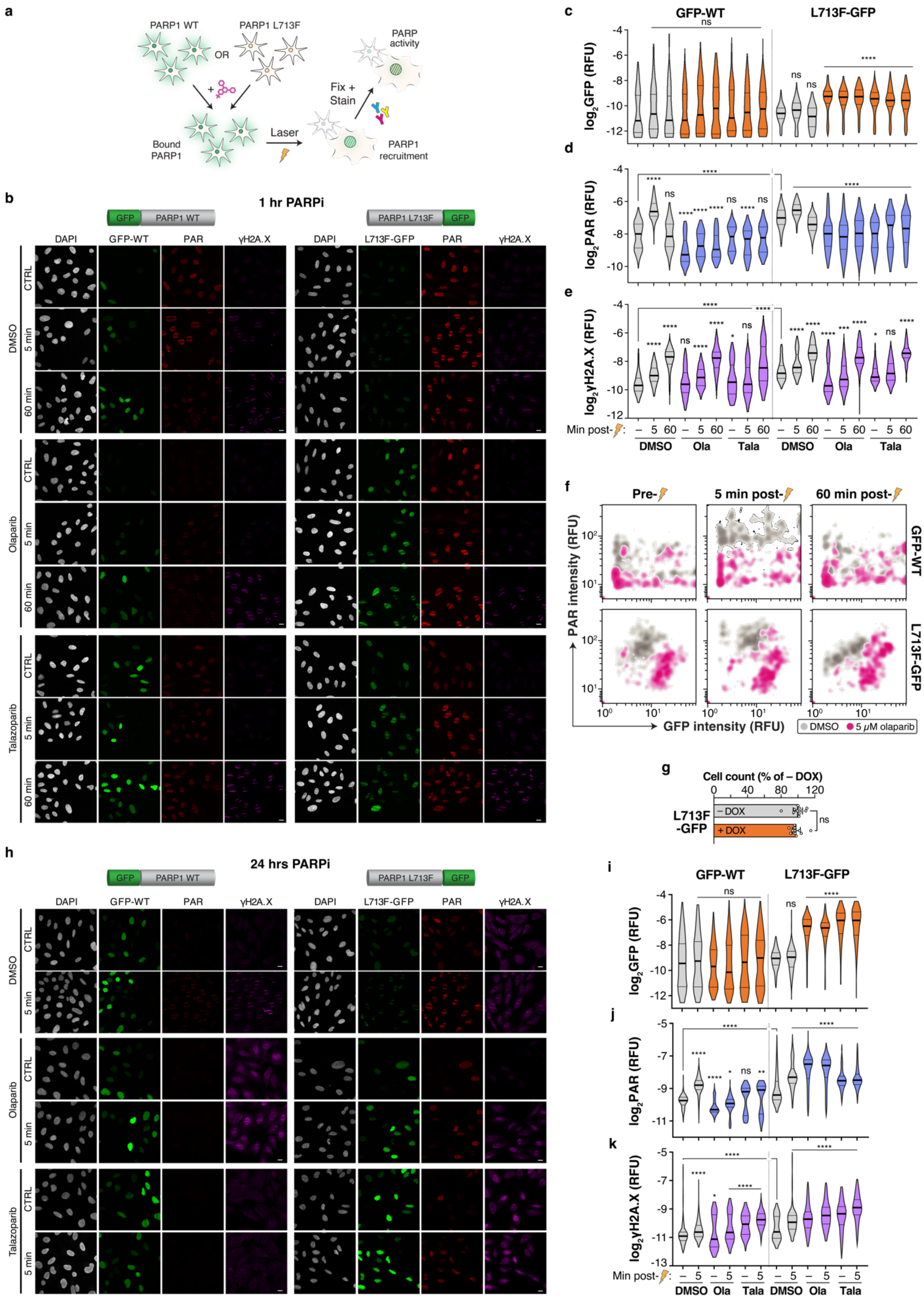
a
PARP1 WT
PARP1 L713F
OR
Bound PARP1
Laser
Fix + Stain
PARP activity
PARP1 recruitment
b
1 hr PARPi
GFP
PARP1 WT
PARP1 L713F
DAPI
GFP-WT
PAR
γH2A.X
L713F-GFP
DMSO
Olaparib
Talazoparib
CTRL
5 min
60 min
c
GFP-WT
L713F-GFP
log2GFP (RFU)
d
log2PAR (RFU)
e
log2γH2A.X (RFU)
Min post-
DMSO
Ola
Tala
f
Pre-
5 min post-
60 min post-
PAR intensity (RFU)
GFP intensity (RFU)
DMSO
5 μM olaparib
g
Cell count (% of – DOX)
L713F -GFP
– DOX
+ DOX
h
24 hrs PARPi
i
j
k

**Supplementary Figure 9. PARP1 L713F adheres to PARP1 biology in DNA damage repair and response to PARPi. a**, Graphical procedure for monitoring PARP-GFP intensity and activity at DNA damage sites with or without PARPi. U-2 OS GFP-PARP1 WT or PARP1 L713F-GFP cells were induced with DOX and pre-treated with DMSO, 5 μM olaparib, or 0.5 μM talazoparib for 1 or 24 hours before microirradiation and fixation after 5 or 60 minutes. **b**, Representative micrographs of GFP-PARP1 WT, L713F-GFP, PAR, and γH2A.X recruitment dynamics to sites of microirradiation after 1 hour of PARPi. For all micrographs, DAPI – blue; GFP-PARP1 – green; PAR – red; γH2A.X – magenta and scale bar = 20 μm. **c**, Violin plots of $log_2$-transformed GFP, PAR (**d**), and γH2A.X (**e**) intensity following 1-hour PARPi incubation and post-microirradiation (5 and 60 minutes). The medians (thick lines) and quartiles (thin lines) are shown for each plot, while DMSO-treated samples are represented in grey and PARPi-treated counterparts are colored. Data are from 229 total cells per group and compiled from three independent experiments. **f**, Two-dimensional dynamics of WT or L713F PARP1-GFP stabilization and PAR abundance in the absence (grey) or presence (magenta) of olaparib for 1 hour by density plot (corresponding to data in **c** and **d**). **g**, Normalized cell counts from U-2 OS pINDUCER20-PARP1 L713F-GFP #5 cells after 48 hours in the absence or presence of 1 μg/mL doxycycline. Means and 95% confidence intervals are shown. Results are pooled from two independent experiments and comprise of 11 total replicates. **h**, Representative micrographs after 24 hours of PARPi, set up as above. **i**, Violin plots of $log_2$-transformed GFP, PAR (**j**), and γH2A.X (**k**) intensity following 24-hour PARPi incubation and post-microirradiation (5 minutes). Plots are set up identically as above. Data are from 282 total cells (except $n_{L713F\text{-}Tala\text{-}5min}$: 208) and compiled from three independent experiments. ns – not significant; * – $p < 0.05$; *** – $p < 0.001$, **** – $p < 0.0001$ by Kruskal-Wallis analyses with multiple comparisons to pre-irradiated DMSO treatments for each group (Dunn's test; **c**-**e** and **i**-**k**; Kruskal-Wallis statistic = $12.33_{\mathbf{c,WT}}$, $533.8_{\mathbf{c,L713F}}$, $726.7_{\mathbf{d,WT}}$, $572.4_{\mathbf{d,L713F}}$, $652.9_{\mathbf{e,WT}}$, $836.2_{\mathbf{e,L713F}}$, $12.25_{\mathbf{i,WT}}$, $821.7_{\mathbf{i,L713F}}$, $498.9_{\mathbf{j,WT}}$, $352.5_{\mathbf{j,L713F}}$, $300.1_{\mathbf{k,WT}}$, $293.6_{\mathbf{k,L713F}}$), two-tailed Mann-Whitney U test (**d**, **e**, **j**, and **k** [comparison of WT and L713F pre-irradiated DMSO controls], Mann-Whitney U = $7787_{\mathbf{d}}$, $8464_{\mathbf{e}}$, $19116_{\mathbf{j}}$, $26745_{\mathbf{k}}$), or two-tailed t test (**g**; t, df = 0.5787, 20). RFU – relative fluorescence units.

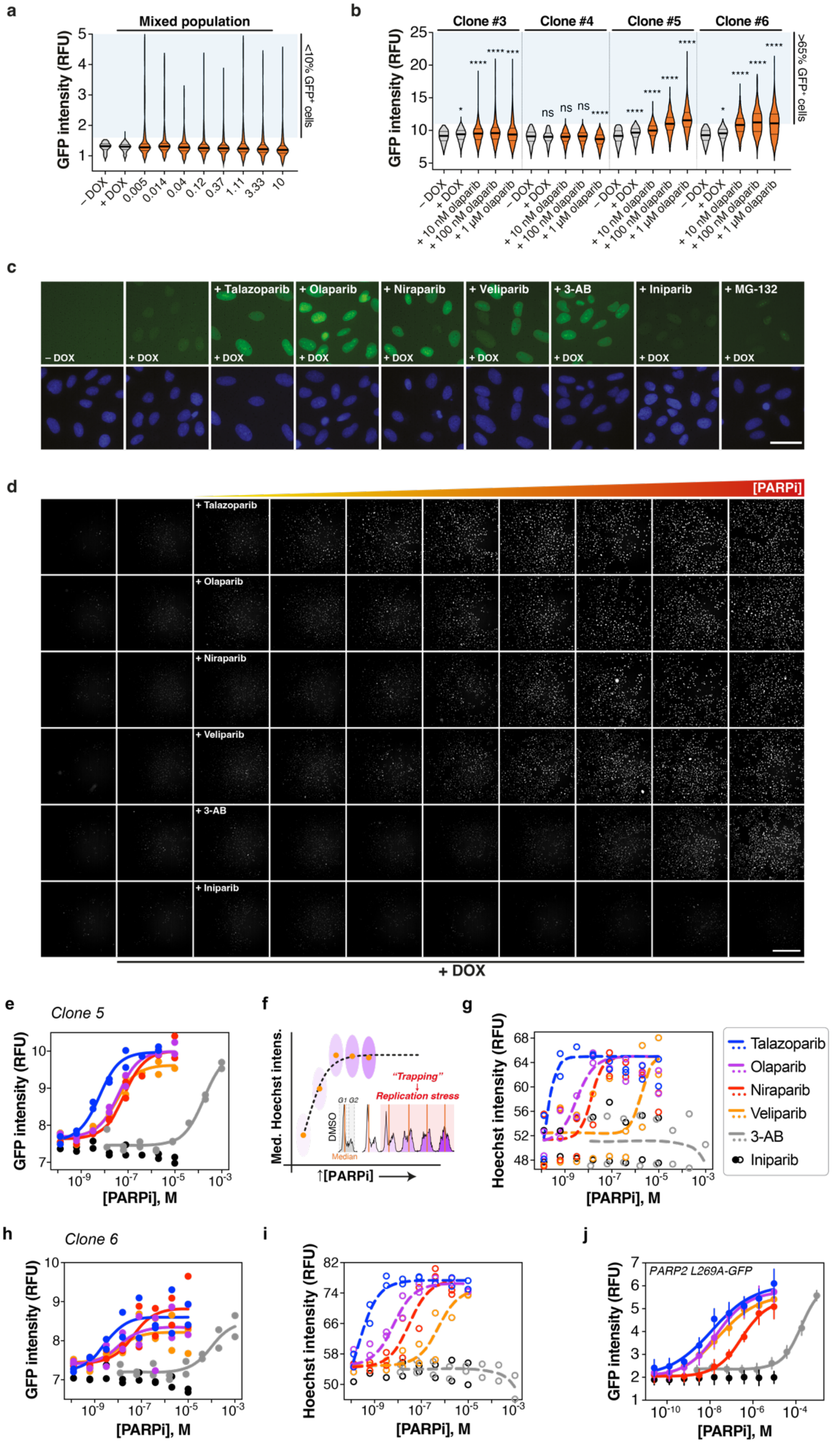

a
Mixed population
GFP intensity (RFU)
<10% GFP+ cells
– DOX
+ DOX
0.005
0.014
0.04
0.12
0.37
1.11
3.33
10
b
Clone #3
Clone #4
Clone #5
Clone #6
GFP intensity (RFU)
>65% GFP+ cells
– DOX
+ DOX
+ 10 nM olaparib
+ 100 nM olaparib
+ 1 μM olaparib
c
+ Talazoparib
+ Olaparib
+ Niraparib
+ Veliparib
+ 3-AB
+ Iniparib
+ MG-132
d
[PARPi]
+ DOX
e
Clone 5
GFP intensity (RFU)
[PARPi], M
f
Med. Hoechst intens.
"Trapping"
Replication stress
DMSO
Median
[PARPi]
g
Hoechst intensity (RFU)
[PARPi], M
Talazoparib
Olaparib
Niraparib
Veliparib
3-AB
Iniparib
h
Clone 6
GFP intensity (RFU)
[PARPi], M
i
Hoechst intensity (RFU)
[PARPi], M
j
PARP2 L269A-GFP
GFP intensity (RFU)
[PARPi], M

**Supplementary Figure 10. Clonal characterization of live PARP1 L713F-GFP-expressing cells (related to Figure 5). a**, Representative violin plots of live-cell GFP intensity from pooled ("mixed population") U-2 OS PARP1 L713F-GFP cells after 24 hours of a talazoparib gradient. Measurements are of 501 cells per condition. **b**, Representative violin plots of live-cell GFP intensity from clonal U-2 OS PARP1 L713F-GFP cell populations after 24 hours of an olaparib gradient. Clonal populations of U-2 OS PARP1 L713F-GFP cells were incubated with olaparib at various concentrations for 24 hours. Measurements are of 499 cells per condition. ns – not significant, * – $p < 0.05$, *** – $p < 0.001$, **** – $p < 0.0001$ by Kruskal-Wallis test with comparisons to DMSO treatments (+DOX) for each group (Dunn's test; Kruskal-Wallis statistic = $40.35_{clone3}$, $60.47_{clone4}$, $817.5_{5clone5}$, $620.0_{clone6}$). For violin plots in **a** and **b**, medians (thick lines) and quartiles (thin lines) are shown, while DMSO-treated samples are represented in grey and PARPi-treated counterparts are orange. Blue regions are a visual reference for population shifts. **c**, Representative live-cell GFP and Hoechst micrographs of clone 5 cells induced by DOX and incubated with 10 µM talazoparib, 10 µM olaparib, 10 µM niraparib, 10 µM veliparib, 1000 µM 3-aminobenzamide (3-AB), or 10 µM iniparib for 24 hours. 5 µM MG-132 for 6 hours was used as a control. Scale bar = 50 µm. **d**, Representative live-cell GFP micrographs (greyscale) from a dose-response of talazoparib, olaparib, niraparib, veliparib, 3-AB, and iniparib of induced clone 5 cells after 24 hours. Scale bar = 500 µm. **e**, Summary median GFP intensity saturation curves from clone 5 cells. Mean data points from two independent experiments and respective curve fittings are shown. **f**, Graphic describing the use of median Hoechst intensity (DNA content) as a readout of PARPi-induced DNA replication stress due to PARP trapping. **g**, Summary Hoechst intensity (DNA content) saturation curves from clone 5 cells. Mean data points from two independent experiments and respective curve fittings are shown. **h**, Summary median GFP intensity and Hoechst intensity (DNA content, **i**) saturation curves from clone 6 cells. Mean data points from two independent experiments and respective curve fittings are shown. **j**, Summary median GFP intensity saturation curves with U-2 OS cells constitutively expressing PARP2 L269A-GFP cells (extended dataset related to **Figure 2i**). Means from five independent experiments ± SEM and respective curve fittings are shown. RFU – relative fluorescence units.

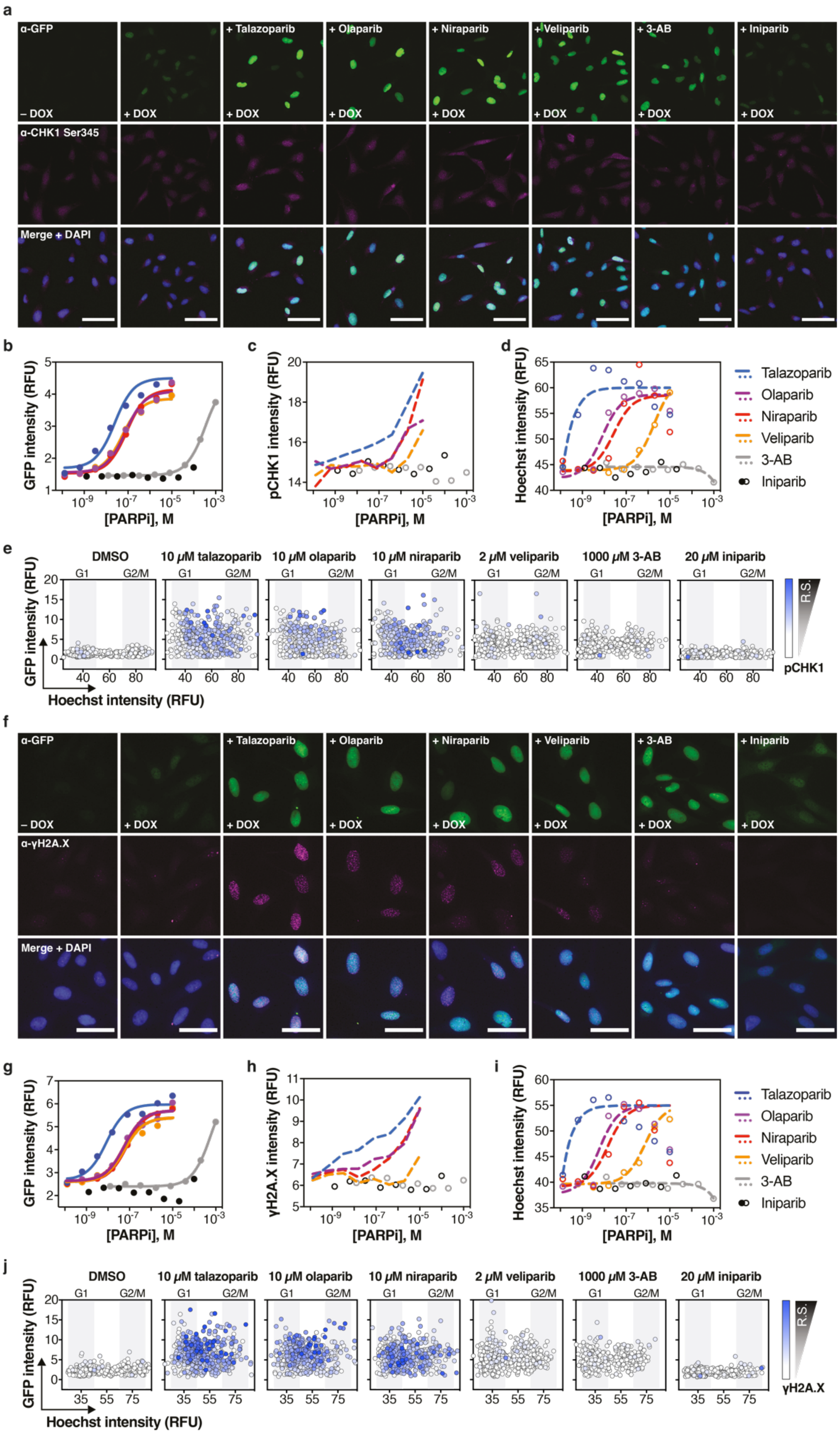
a
α-GFP
– DOX
+ DOX
+ Talazoparib
+ Olaparib
+ Niraparib
+ Veliparib
+ 3-AB
+ Iniparib
α-CHK1 Ser345
Merge + DAPI
b
GFP intensity (RFU)
[PARPi], M
c
pCHK1 intensity (RFU)
d
Hoechst intensity (RFU)
Talazoparib
Olaparib
Niraparib
Veliparib
3-AB
Iniparib
e
DMSO
10 μM talazoparib
10 μM olaparib
10 μM niraparib
2 μM veliparib
1000 μM 3-AB
20 μM iniparib
G1
G2/M
R.S.
pCHK1
Hoechst intensity (RFU)
f
α-γH2A.X
g
h
γH2A.X intensity (RFU)
i
j
γH2A.X

**Supplementary Figure 11. Multiparametric interrogation of replication stress in fixed PARP1 L713F-GFP-expressing cells exposed to PARP inhibitors.** **a**, Representative micrographs of induced U-2 OS PARP1 L713F-GFP clone 5 cells incubated with increasing concentrations of talazoparib, olaparib, niraparib, veliparib, iniparib, or 3-AB for 24 hours and stained for GFP (target engagement), CHK1 Ser345 (pCHK1; replication stress/DNA damage), and Hoechst 33342 (DNA content; in merged image). Scale bar = 200 µm. Median GFP (**b**), CHK1 Ser345 (**c**), and Hoechst (**d**) intensities. **e**, Per-cell, three-dimensional visualization of GFP (y-axis), Hoechst (x-axis), and CHK1 Ser345 (blue gradient) after DMSO, 10 µM talazoparib, 10 µM olaparib, 10 µM niraparib, 2 µM veliparib, 1000 µM 3-AB or 20 µM iniparib. Estimated G1 and G2/M cell populations are demarcated. n = 500 cells shown per condition. **f**, Representative micrographs of induced clone 5 cells incubated with increasing concentrations of talazoparib, olaparib, niraparib, veliparib, iniparib, or 3-AB for 24 hours and stained for GFP, γH2A.X (replication stress/DNA damage), and Hoechst 33342 (in merged image). Scale bar = 50 µm. Median GFP (**g**), γH2A.X (**h**), and Hoechst (**i**) intensities. **j**, Per-cell three-dimensional visualization of GFP (y-axis), Hoechst (x-axis), and γH2A.X (blue gradient) after DMSO, 10 µM talazoparib, 10 µM olaparib, 10 µM niraparib, 2 µM veliparib, 1000 µM 3-AB or 20 µM iniparib. Estimated G1 and G2/M cell populations are demarcated. 500 cells per condition are shown. In all cases, representative experiments and lines of best fit are shown; RFU – relative fluorescence units and blue gradient – increasing replication stress/DNA damage (CHK1 Ser345 or γH2A.X; R.S.).

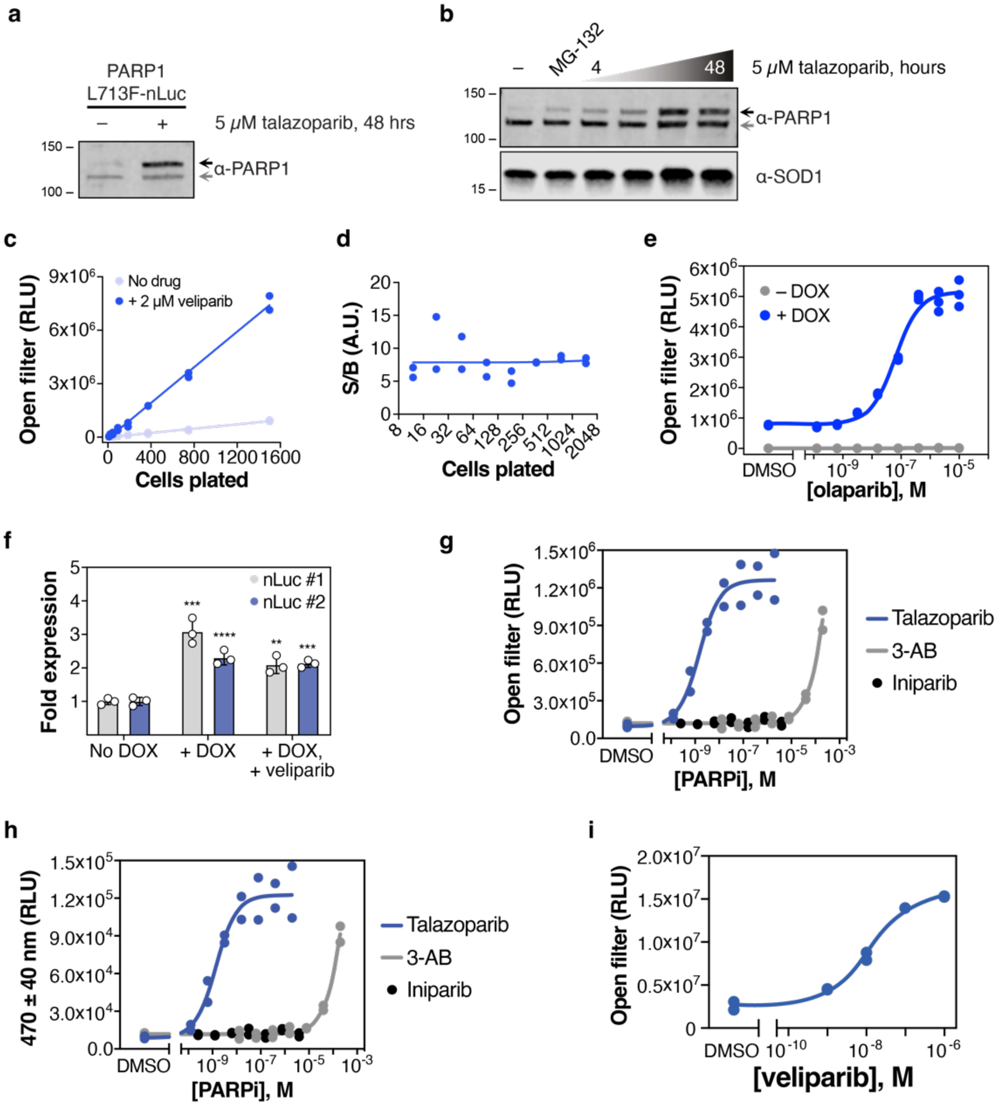
a
PARP1
L713F-nLuc
5 μM talazoparib, 48 hrs
α-PARP1
b
MG-132
5 μM talazoparib, hours
α-PARP1
α-SOD1
c
Open filter (RLU)
No drug
+ 2 μM veliparib
Cells plated
d
S/B (A.U.)
Cells plated
e
Open filter (RLU)
– DOX
+ DOX
[olaparib], M
f
Fold expression
nLuc #1
nLuc #2
No DOX
+ DOX
+ DOX, + veliparib
g
Open filter (RLU)
Talazoparib
3-AB
Iniparib
[PARPi], M
h
470 ± 40 nm (RLU)
Talazoparib
3-AB
Iniparib
[PARPi], M
i
Open filter (RLU)
[veliparib], M

**Supplementary Figure 12. Assessment of PARP1 L713F-nLuc stabilization by PARPi (related to Figure 6). a**, U-2 OS PARP1 L713F-nLuc #3 cells pre-treated with 1 μg/mL DOX were exposed to DMSO or 5 μM talazoparib for 48 hours. **b**, A representative blot (n=2) of cells treated with 5 μM talazoparib for 4-48 hours (as in **a**). As controls, DMSO was added for 48 hours and 5 μM MG-132 for 6 hours. For both, black arrow – PARP1 L713F-nLuc; grey arrow – endogenous PARP1. **c**, Cells pre-treated as in **a**, plated at different densities, and treated with DMSO (No drug, light blue) or 2 μM veliparib (blue) for 24 hours before luminescence readings. Linear regressions are shown for both data sets. Data points from two independent experiments are shown. **d**, Signal-to-background calculations across different cell numbers plated (from **c**). Linear regression illustrates the consistency of the S/B. Data points from two independent experiments are shown. **e**, Cells (as in **a**) were plated in the absence (grey) or presence of 1 μg/mL DOX (blue) were exposed to DMSO or serial dilutions of olaparib for 24 hours prior to luminescence detection. Data points from three independent experiments and relevant curve fitting are shown. **f**, Fold expression of nLuc in pooled HCT116 pCW57.1-PARP1 L713F-nLuc cells after normalization to GAPDH and β-actin by RT-qPCR. Means ± SD are shown for a representative experiment. ** – $p < 0.01$; *** – $p < 0.001$; **** – $p < 0.0001$ by two-way ANOVA with multiple comparisons to "No DOX" for each primer pair (Dunnett's test, F [DFn, DFd]: $F_{Interaction}$ [2, 12] = 6.517, $F_{Row\ Factor}$ [1, 12] = 5.544, $F_{Column\ Factor}$ [2, 12] = 89.70). **g**, After induction, HCT116 pCW57.1-PARP1 L713F-nLuc cells were treated with serial dilutions of talazoparib, 3-AB, or iniparib diluted in Fluorobrite DMEM for 24 hours before luminescence detection on an open filter setting. Individual data points and relevant curve fittings are shown for two independent experiments. **i**, The same experiments from **h** but measured on a nLuc-specific filter. Individual data points and relevant curve fittings are shown. **j**, HCT116 pLenti CMV Blast-PARP1 L713F-nLuc cells were treated with DMSO or veliparib for 24 hours before measuring luminescence on an open filter. Individual data points and relevant curve fittings are shown for two independent experiments. RLU – relative luminescence units.

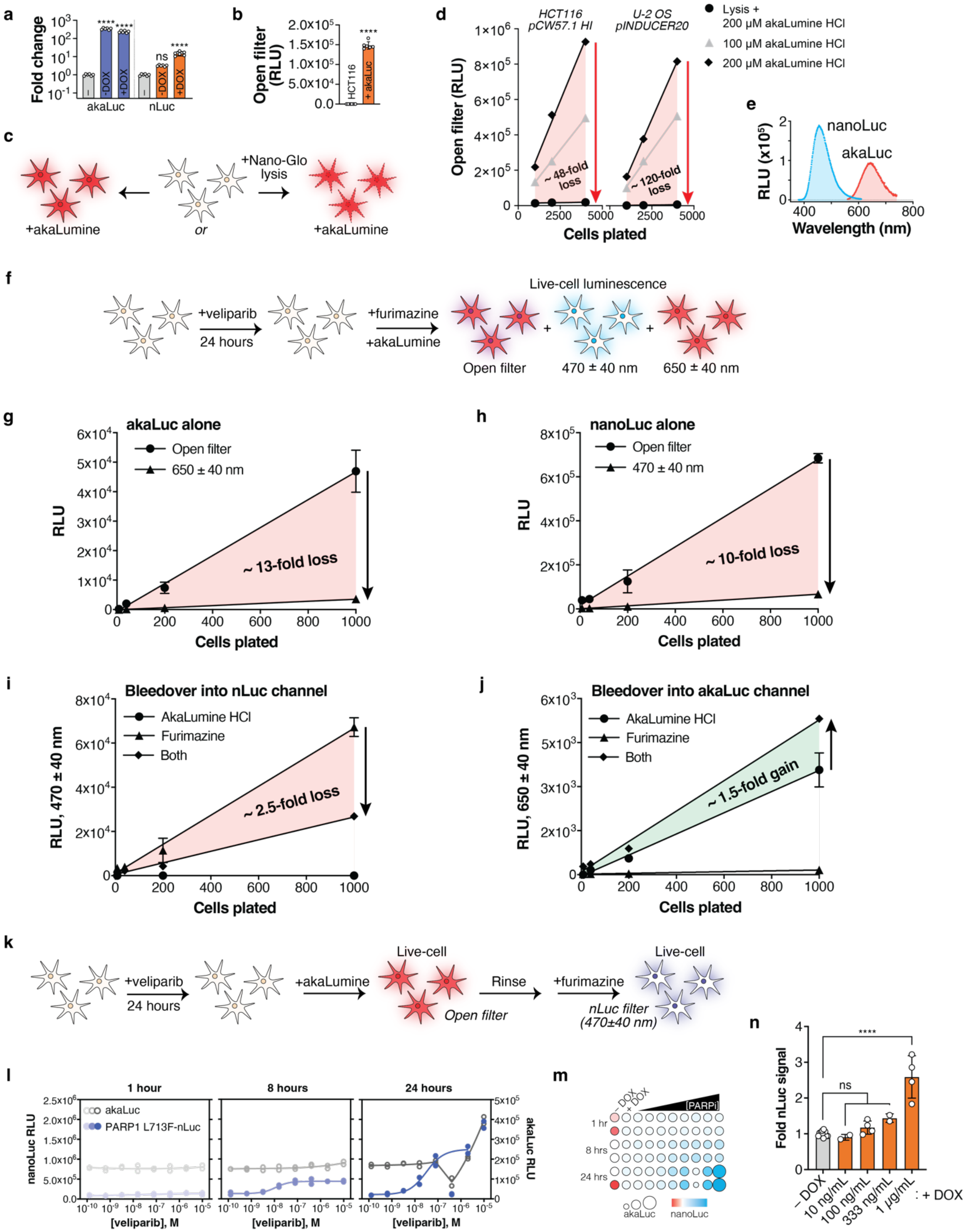

a
Fold change
akaLuc
nLuc
-DOX
+DOX
ns
b
Open filter (RLU)
HCT116
+ akaLuc
c
+Nano-Glo lysis
+akaLumine
or
d
HCT116 pCW57.1 HI
U-2 OS pINDUCER20
Lysis + 200 µM akaLumine HCl
100 µM akaLumine HCl
200 µM akaLumine HCl
Open filter (RLU)
~ 48-fold loss
~ 120-fold loss
Cells plated
e
RLU (x10⁵)
nanoLuc
akaLuc
Wavelength (nm)
f
+veliparib
24 hours
+furimazine
+akaLumine
Live-cell luminescence
Open filter
470 ± 40 nm
650 ± 40 nm
g
akaLuc alone
Open filter
650 ± 40 nm
~ 13-fold loss
RLU
Cells plated
h
nanoLuc alone
Open filter
470 ± 40 nm
~ 10-fold loss
RLU
Cells plated
i
Bleedover into nLuc channel
AkaLumine HCl
Furimazine
Both
~ 2.5-fold loss
RLU, 470 ± 40 nm
Cells plated
j
Bleedover into akaLuc channel
AkaLumine HCl
Furimazine
Both
~ 1.5-fold gain
RLU, 650 ± 40 nm
Cells plated
k
+veliparib
24 hours
+akaLumine
Live-cell
Open filter
Rinse
+furimazine
Live-cell
nLuc filter (470±40 nm)
l
1 hour
8 hours
24 hours
akaLuc
PARP1 L713F-nLuc
nanoLuc RLU
akaLuc RLU
[veliparib], M
m
– DOX
+ DOX
[PARPi]
1 hr
8 hrs
24 hrs
akaLuc
nanoLuc
n
Fold nLuc signal
****
ns
– DOX
10 ng/mL
100 ng/mL
333 ng/mL
1 µg/mL
: + DOX

**Supplementary Figure 13. AkaLuc suitability as an nLuc normalization factor (related to Figure 6). a**, RT-qPCR analysis of parental HCT116 (–; grey) or HCT116 pCW57.1-PARP1 L713F-nLuc/pLenti CMV Blast-akaLuc cells treated with -/+ DOX for 24 hours. akaLuc (blue) and nLuc (orange) fold expression was determined by normalization to GAPDH and β-actin and plotted in $\log_{10}$ scale. Means of two independent experiments in triplicate ± SD are shown. **b**, Cells were treated as in **a**, then incubated with 200 µM akaLumine HCl immediately before luminescence measurement on an open filter. Means of two independent experiments in triplicate ± SD are shown. **c**, Graphical procedure for determining akaLuc amenability to lytic conditions. **d**, Induced HCT116 pCW57.1-PARP1 L713F-nLuc/pLenti CMV Blast-akaLuc or U-2 OS pINDUCER20-PARP1 L713F-nLuc/pLenti-CMV-blast-akaLuc cells plated at different densities were incubated with 200 µM akaLumine HCl following Nano-Glo lysis reagent or at 100 or 200 µM in FluoroBrite DMEM with live cells immediately before luminescence readings. A single experiment and estimated luminescence intensity change due to procedural variations are shown for each. **e**, Emission and intensity spectra of L713F-nLuc (blue) and akaLuc (red). **f**, Graphical procedure of concomitant measuring of nLuc and akaLuc signals following 1 µM veliparib treatment in live HCT116 pCW57.1-PARP1 L713F-nLuc/pLenti CMV Blast-akaLuc cells. **g**, Comparison of akaLuc detection using an open filter or akaLuc-specific filter, as in **f**. Means of two independent experiments ± SD and estimated luminescence intensity change due to procedural variations are shown. **h**, Comparison of L713F-nLuc detection using an open filter or nLuc-specific filter, as in **f**. Means of two independent experiments ± SD and estimated luminescence intensity change due to procedural variations are shown. **i**, Comparison of nLuc filter signal when cells are incubated with akaLumine HCl, furimazine, or both simultaneously at different plating numbers. Means of two independent experiments ± SD and estimated luminescence intensity change due to procedural variations are shown. **j**, Comparison of akaLuc filter signal, as in **i**. Means of two independent experiments ± SD and estimated luminescence intensity change due to procedural variations are shown. **k**, Graphical procedure for sequential measurement of akaLuc followed by L713F-nLuc in live cells. **l**, Time-dependent maturation of L713F-nLuc signal (left axis, blue) in relation to akaLuc signal (right axis, grey) in response to a veliparib gradient. A single experiment performed in duplicate is shown. **m**, Normalization matrix relating L713F-nLuc signal to akaLuc signal with data from **l**. Circle size – akaLuc intensity; red-white-blue gradient – nLuc intensity. **n**, Cells were incubated with the indicated concentration of doxycycline (DOX) for 24 hours before sequential measurement of akaLuc and nLuc bioluminescence. nLuc signals are normalized to akaLuc and represented as fold change relative to "– DOX". Individual replicates are indicated from two independent experiments ± SD (subset of data from **Fig. 6c**). In all cases, ns – not significant; **** – $p < 0.0001$ by one-way ANOVA analysis with multiple comparisons (Dunnett's test; to parental HCT116 cells for each primer set [**a**; $F_{\text{Treatment akaLuc}}$ (DFn, DFd) = 237.3 (2, 15), $F_{\text{Treatment nLuc}}$ (DFn, DFd) = 76.85 (2, 15)] or – DOX [**n;** $F_{\text{Treatment}}$ (DFn, DFd) = 31.31 (4, 19)]) or two-tailed t test (**b**; t, df = 38.79, 10). RLU – relative luminescence units.

**a**

| Cell line/vector | Z' | S/B |
|---|---|---|
| HCT116 pCW57.1 | 0.51 | 11.19 |
| U-2 OS pINDUCER | 0.77 | 6.47 |
| HCT116 pLenti | 0.83 | 5.88 |

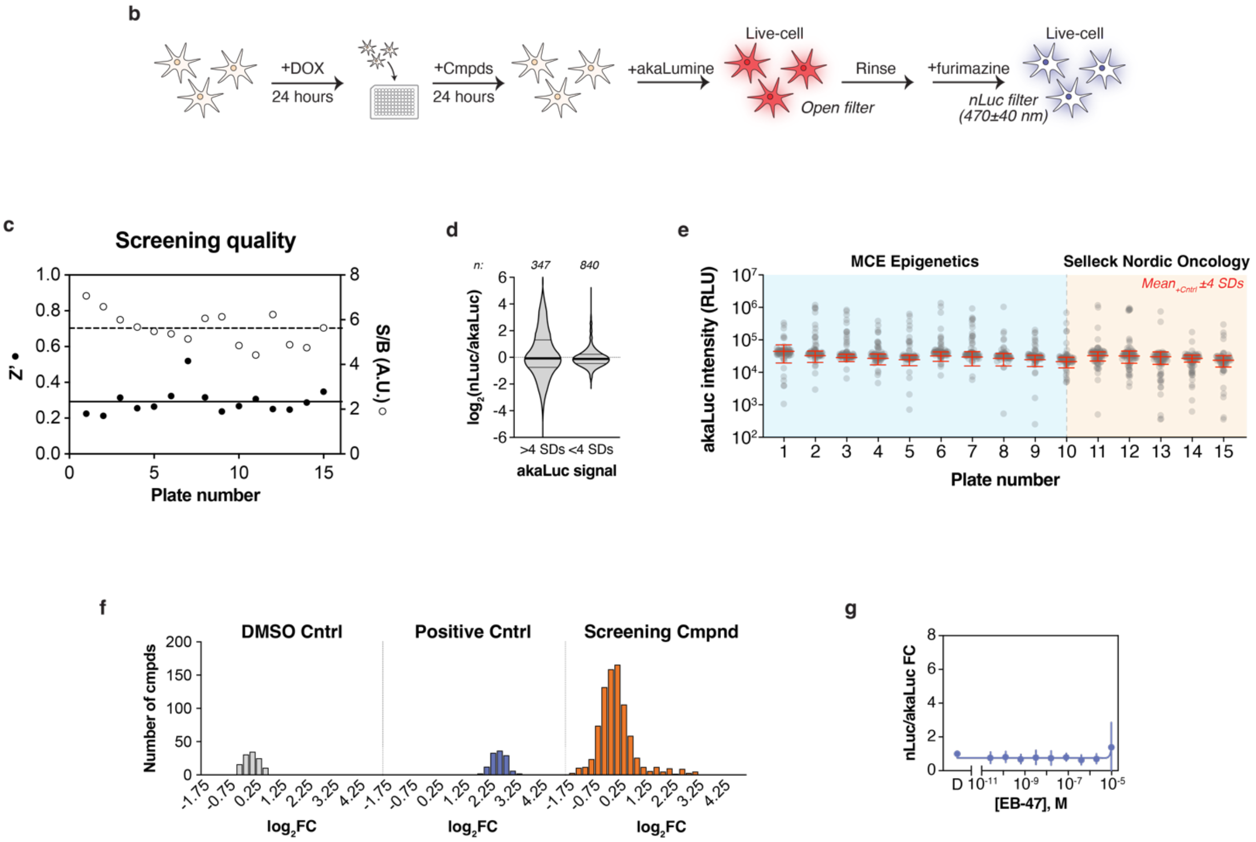

**Supplementary Figure 14. Data quality and analysis from PARP1 L713F biophysical perturbagen screen (related to Figure 6). a**, A comparison of Z' and signal-to-background (S/B) values across cell and expression systems from earlier testing. $Z' = 1 - [3\sigma_+ + 3\sigma_-] / |\mu_+ - \mu_-|$ and S/B = [mean signal] / [mean background], as defined by Zhang, Chung, & Oldenburg[6]. **b**, A graphical depiction of the screening procedure. **c**, Screening quality summary on a per plate basis looking at Z' (left axis, closed circles) and S/B (right axis, open circles) based on the positive and negative controls on each plate. **d**, Violin plots of $log_2$-transformed nLuc/akaLuc ratios for the screening library triaged by akaLuc signal being within (right) or beyond four SDs (left) of controls. Medians (thick lines) and quartiles (thin lines), as well as the number of screening compounds falling into each category are shown. **e**, Per-plate scatter plots of akaLuc intensity from screening compounds with overlayed means ± 4 SDs from positive controls (red). Library designations are also labeled (MCE Epigenetics – blue; Selleck Nordic Oncology – orange). **f**, Histograms showing improved normality of $log_2$-transformed nLuc/akaLuc ratios for DMSO control (negative; grey), veliparib control (positive; blue), and screening library compounds (orange). Bin width 0.25 **g**, Follow-up dose-response analysis with L713F-nLuc and EB-47. Means from three independent experiments (in duplicate) ± SD and relevant curve fittings are shown. A.U. – arbitrary unit, RLU – relative luminescence unit, FC – fold change.

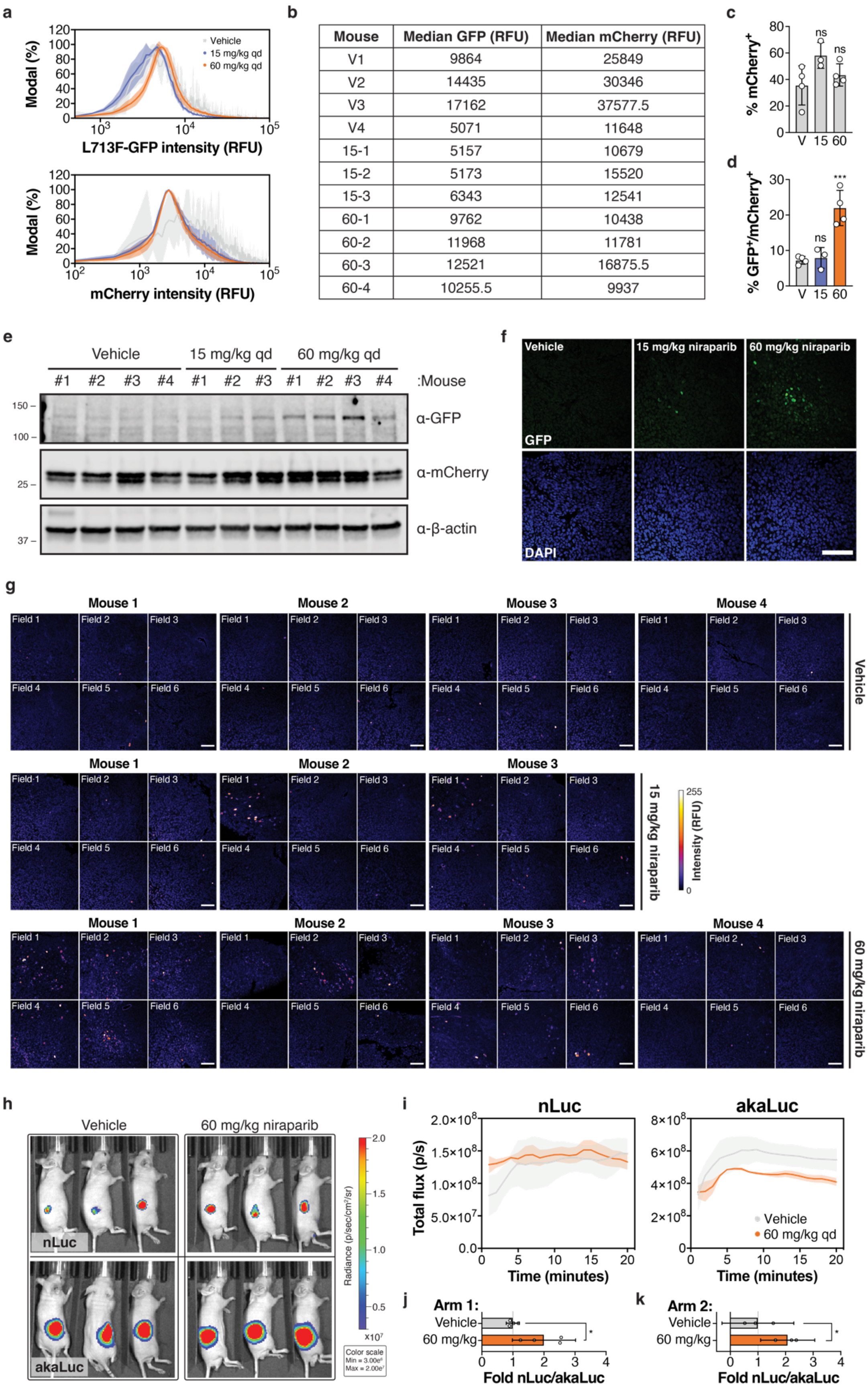

| Mouse | Median GFP (RFU) | Median mCherry (RFU) |
|---|---|---|
| V1 | 9864 | 25849 |
| V2 | 14435 | 30346 |
| V3 | 17162 | 37577.5 |
| V4 | 5071 | 11648 |
| 15-1 | 5157 | 10679 |
| 15-2 | 5173 | 15520 |
| 15-3 | 6343 | 12541 |
| 60-1 | 9762 | 10438 |
| 60-2 | 11968 | 11781 |
| 60-3 | 12521 | 16875.5 |
| 60-4 | 10255.5 | 9937 |

**Supplementary Figure 15. Supporting data for *in vivo* CeTEAM analyses (related to Figures 7 and 8).** **a**, Average L713F-GFP and mCherry intensity histograms from flow cytometry of live vehicle- (grey), 15 mg/kg- (blue), and 60 mg/kg-treated (orange) HCT116 tumor cells. Histograms are normalized to the mode for each dataset. The error bands denote the data ranges from $n_{Vehicle} = 4$, $n_{15mg/kg} = 3$, and $n_{60mg/kg} = 4$ mice. **b**, Median L713F-GFP and mCherry fluorescence intensities from tumor cells by flow cytometry used to calculate fold change of PARPi target engagement. **c**, Percent of live tumor cells identified as mCherry positive (mCherry$^+$). Means from individual mice ± SD are shown. **d**, Percent of live tumor cells identified as GFP/mCherry double positive (GFP$^+$/mCherry$^+$). Means from individual mice ± SD are shown. ns – not significant, *** – $p < 0.001$ by ordinary one-way ANOVA with multiple comparisons to the DMSO control (Dunnett's test; $F_{Treatment\ (c)}$ [DFn, DFd] = 0.4597 [2, 8], $F_{Treatment\ (d)}$ [DFn, DFd] = 2.821 [2, 8]). **e**, Full western blot from **Fig. 7g**, including mCherry and β-actin loading controls. **f**, anti-GFP staining of representative PARP1 L713F-GFP/mCherry tumor sections from vehicle- (m1, f6), 15 mg/kg- (m7, f5), or 60 mg/kg niraparib-treated mice (m9, f5) counterstained with DAPI. Scale bar = 100 µm. **e**, Micrographs of L713F-GFP saturation in tumor sections by fire LUT. Six fields were taken per section. **f**, Representative bioluminescence overlays for L713F-nLuc and akaLuc radiance from experimental arm 2 (n = 3 mice per group). **g**, Change in total luminescent flux (photons/second) over 20 minutes for both L713F-nLuc and akaLuc following substrate injection in mice treated with vehicle (grey) or 60 mg/kg niraparib (orange). n = 3 mice per group (arm 2); error bands denote the SEM. **h**, Comparison of the mean fold change in L713F-nLuc/akaLuc signal from experimental arm 1 (n = 4 mice per group) and arm 2 (**i**, n = 3 mice per group). In **h** and **i**, error denotes 95% confidence interval, and * – $p < 0.05$ by two-tailed t test (t, df = $[3.064, 6]_{\mathbf{h}}$, $[2.863, 4]_{\mathbf{i}}$). RFU – relative fluorescence units.

a

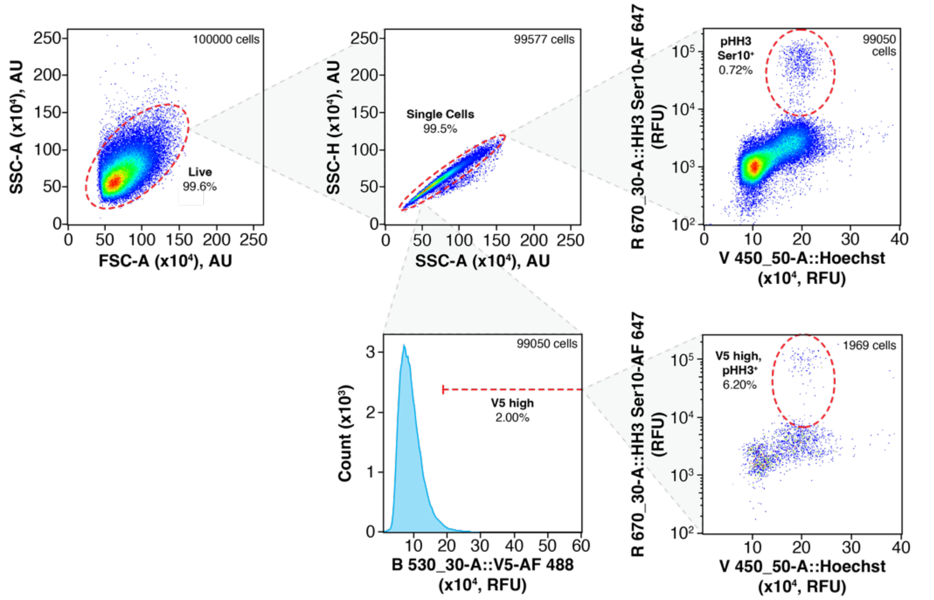

b

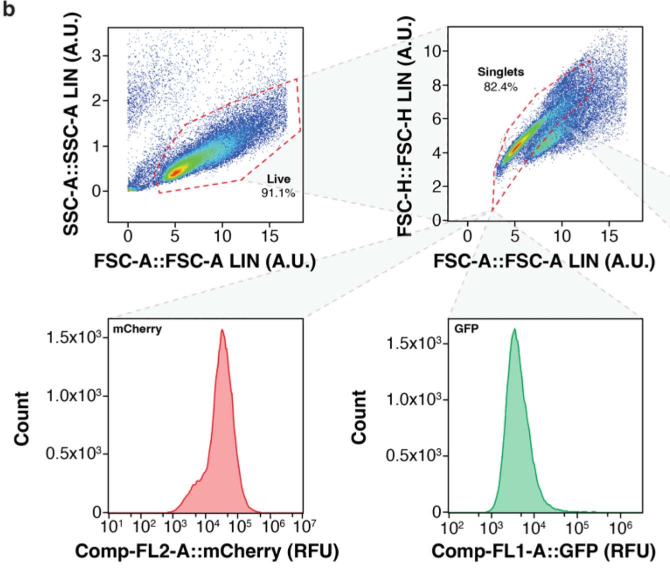

c

| Channel into: | FL1-A GFP | FL2-A mCherry | FL3-A | FL4-A |
|---|---|---|---|---|
| FL1-A GFP | 100% | 2% | 0% | 0% |
| FL2-A mCherry | 2% | 100% | 0% | 0% |
| FL3-A | 0% | 0% | 100% | 0% |
| FL4-A | 0% | 0% | 0% | 100% |

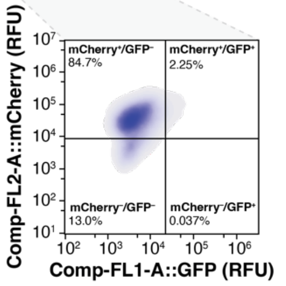

**Supplementary Figure 16. Gating strategies used for flow cytometry analyses. a**, Gating of V5-MTH1 G48E cells. “Live cells” were identified and gated to identify “Single Cells” before identification of “pHH3 Ser10$^{+}$” cells. “Single cells” were gated to identify “V5 high” cells (arbitrarily defined as top 2% of V5 signal in the DMSO control), from which “V5 high/pHH3$^{+}$” cells were identified. **b**, PARP L713F GFP/mCherry gating strategy. “Live cells” were identified and gated as “singlet” cells before identification of “mCherry$^{+}$” and “GFP$^{+}$” cells. This population was also plotted in two dimensions to simultaneously follow mCherry and GFP signals and to establish compensation parameters (**c**). The examples shown are for DMSO-treated cells but applies identically to cells incubated with inhibitors *in vitro* or *in vivo*. Axes are denoted with relevant laser, filter, marker, and fluorescent label. RFU – relative fluorescence units.
